# RIM-BP2 regulates Ca^2+^ channel abundance and neurotransmitter release at hippocampal mossy fiber terminals

**DOI:** 10.1101/2022.08.29.505728

**Authors:** Rinako Miyano, Hirokazu Sakamoto, Kenzo Hirose, Takeshi Sakaba

## Abstract

Synaptic vesicles dock and fuse at the presynaptic active zone (AZ), the specialized site for transmitter release. AZ proteins play multiple roles such as recruitment of Ca^2+^ channels as well as synaptic vesicle docking, priming and fusion. However, the precise role of each AZ protein type remains unknown. In order to dissect the role of RIM-BP2 at mammalian cortical synapses having low release probability, we applied direct electrophysiological recording and super-resolution imaging to hippocampal mossy fiber terminals of RIM-BP2 KO mice. By using direct presynaptic recording, we found the reduced Ca^2+^ currents. The measurements of EPSCs and presynaptic capacitance suggested that the initial release probability was lowered because of the reduced Ca^2+^ influx and impaired fusion competence in RIM-BP2 KO. Nevertheless, larger Ca^2+^ influx restored release partially. Consistent with presynaptic recording, STED microscopy suggested less abundance of P/Q-type Ca^2+^ channels at AZs deficient in RIM-BP2. Our results suggest that the RIM-BP2 regulates both Ca^2+^ channel abundance and transmitter release at mossy fiber synapses.

## Introduction

At chemical synapses, an action potential arriving at the presynaptic terminal opens voltage-gated Ca^2+^ channels, and Ca^2+^ influx through voltage-gated Ca^2+^ channels triggers neurotransmitter release from synaptic vesicles within milliseconds. Synaptic vesicle fusion occurs at a specialized region of the presynaptic membrane called the active zone (AZ), where Ca^2+^ channel clusters and vesicle fusion sites are accommodated via AZ scaffold proteins (Südhof, 2012). For rapid transmission, accumulation of Ca^2+^ channels, co-localization of Ca^2+^ channels and fusion sites, and molecular priming of synaptic vesicles for fusion are required, and the differences among these factors might provide a basis for synaptic diversity. However, it remains unknown how these factors are regulated by AZ-scaffold proteins.

Rab3-interacting molecule-binding proteins (RIM-BPs) represent one principal, conserved family of AZ proteins and bind to RIMs and Ca^2+^ channels (Wang et al., 2000; Hibino et al., 2002). RIM-BP family includes RIM-BP1, RIM-BP2, and RIM-BP3 in mammals, but RIM-BP3 expression is low in the nervous system (Mittelstaedt and Schoch, 2007). In *Drosophila* neuromuscular junctions (NMJs), loss of RIM-BPs decreases Ca^2+^ channel density and reduces release probability (Liu et al., 2011). Moreover, RIM-BP in *Drosophila* NMJs is necessary for tight coupling of synaptic vesicles to Ca^2+^ channels and replenishment of high release probability vesicles (Liu et al., 2011; Müller et al., 2015; Petzoldt et al., 2020). In mammalian synapses, the observations from KO mice are diverse. In RIM-BP1,2 DKO mice, the coupling between Ca^2+^ channels and synaptic vesicles becomes loose, and action potential-evoked neurotransmitter release is reduced at the calyx of Held synapse (Acuna et al., 2015; Butola et al., 2021). At hippocampal CA3-CA1 synapses, RIM-BP2 deletion alters Ca^2+^ channel localization at the AZs without altering total Ca^2+^ influx. Besides, RIM-BP1,2 DKO has no additional effect, indicating that RIM-BP2 dominates the function of RIM-BP isoforms (Grauel et al., 2016; see also Krinner et al., 2021). Tight coupling between Ca^2+^ channels and vesicles is crucial for rapid and efficient transmitter release (Wadel et al., 2007).

In contrast, hippocampal mossy fiber-CA3 synapses are characterized by low release probability (Vyleta and Jonas, 2014). At hippocampal mossy fiber synapses, gSTED imaging suggests that RIM-BP2 KO alters the Munc13-1 cluster number and distribution. In addition, RIM-BP2 deletion decreases the number of docked vesicles (Brockmann et al., 2019). However, because of technical difficulty, direct and quantitative measurements of exocytosis in RIM-BP2 KO terminals have not been performed so far.

To quantify the RIM-BP2 function in mossy fiber-CA3 synapses, we performed whole-cell patch-clamp recordings from WT and RIM-BP2 KO hippocampal mossy fiber boutons. From electrophysiological recordings, we found that the reductions of Ca^2+^ current amplitudes in RIM-BP2 KO terminals. The EPSC and capacitance measurements suggested that the reduction of Ca^2+^ currents was responsible, in part, for that of transmitter release. In addition, they suggested that fusion competence was impaired, which could be overcome by larger Ca^2+^ influx into the terminal. Using STED imaging, we found the abundance of P/Q-type Ca^2+^ channels in terminals to be reduced in RIM-BP2 KO. We suggest that RIM-BP2 regulates the number of P/Q-type Ca^2+^ channels at the AZ and is critical for transmitter release at hippocampal mossy fiber terminals.

## Results

### RIM-BP2 KO reduces presynaptic calcium currents at hippocampal mossy fiber boutons

To investigate the roles of RIM-BP2 in synaptic transmission, we examined the kinetics of exocytosis and Ca^2+^ influx in WT and RIM-BP2 KO synapses using presynaptic whole-cell capacitance measurements (Fig. 1). When the terminal was depolarized from -80 mV to +10 mV for 10 ms, Ca^2+^ currents and membrane capacitance were recorded as shown in Fig. 1A. During the depolarizing pulse, capacitance was not measured due to large conductance changes, and the increase was used for measuring synaptic vesicle exocytosis. To determine the size of the RRP and the time course of exocytosis, we measured capacitance changes (ΔC_m_) in response to various durations of depolarization. Here, the length of the depolarizing pulse was varied between 5 ms and 100 ms, and ΔC_m_ was plotted against pulse duration (Fig. 1B and Figure 1-figure supplement 1, bottom). ΔC_m_ became larger as the duration of pulse was prolonged, but the increase started to be saturated at a 100 ms pulse in WT terminals, suggesting depletion of the RRP. ΔC_m_ in response to a 100 ms pulse was 52 ± 8.1 fF (n = 9, Fig. 1B). The amplitude corresponds to the fusion of about 500-600 synaptic vesicles (Hallermann et al., 2003; Midorikawa and Sakaba, 2017). The time course of ΔC_m_ could be fitted by a single exponential with a time constant of 59 ± 12 ms. In RIM-BP2 KO terminals, ΔC_m_ in response to a 100 ms pulse was somewhat smaller (41 ± 5.9 fF, n = 8) than that of WT terminals, and the release time constant was somewhat slower (64 ± 14 ms) than in WT terminals. We should note that the time constant is not necessarily reliable because the ΔC_m_ values do not reach a plateau value at 100 ms pulse in some terminals. Although the capacitance jumps of KO may be smaller for shorter pulses, capacitance measurements are not sensitive enough to detect small changes (< 5 fF, 50 vesicles, see Fig. 5 & Fig. 6). Nevertheless, the differences in the time course of capacitance were not significantly different between WT and KO (p = 0.23 from ANOVA) and the effects on release examined by strong pulses depleting the RRP was relatively minor. The amplitudes of Ca^2+^ currents were smaller in RIM-BP2 KO terminals as compared to WT terminals (Fig. 1B and Figure 1-figure supplement 1, top, p < 0.02 from ANOVA). The Ca^2+^ current in response to a 100 ms pulse was reduced by ∼30% in RIM-BP2 KO terminals (27 ± 3.9 pA, n = 8) compared to WT terminals (44 ± 5.4 pA, n = 9) (p = 0.0248, t-test, Fig. 1B). Thus, the results show that RIM-BP2 deletion reduces presynaptic Ca^2+^ influx, but effects on the RRP size and the time course of release measured by capacitance measurements are relatively minor.

To investigate the Ca^2+^ influx in more detail, we examined the voltage dependence of Ca^2+^ currents. Fig. 1C shows voltage step protocols and representative current traces. In both genotypes, Ca^2+^ currents started to be activated at around -20 mV, and maximal amplitude was observed at ∼+10 mV (Fig. 1D). There was no major difference in the voltage dependence of Ca^2+^ currents between WT and RIM-BP2 KO mice. Consistently, activation time constants of Ca^2+^ currents were similar between WT and RIM-BP2 KO, when the terminal was depolarized to +10 mV (τ = 1.4 ± 0.2 ms in WT, n = 6; τ = 1.3 ± 0.2 ms in KO, n = 4). These results suggest a reduction in the number of Ca^2+^ channels rather than changes in activation kinetics is responsible for the current decrease observed.

**Figure 1.**
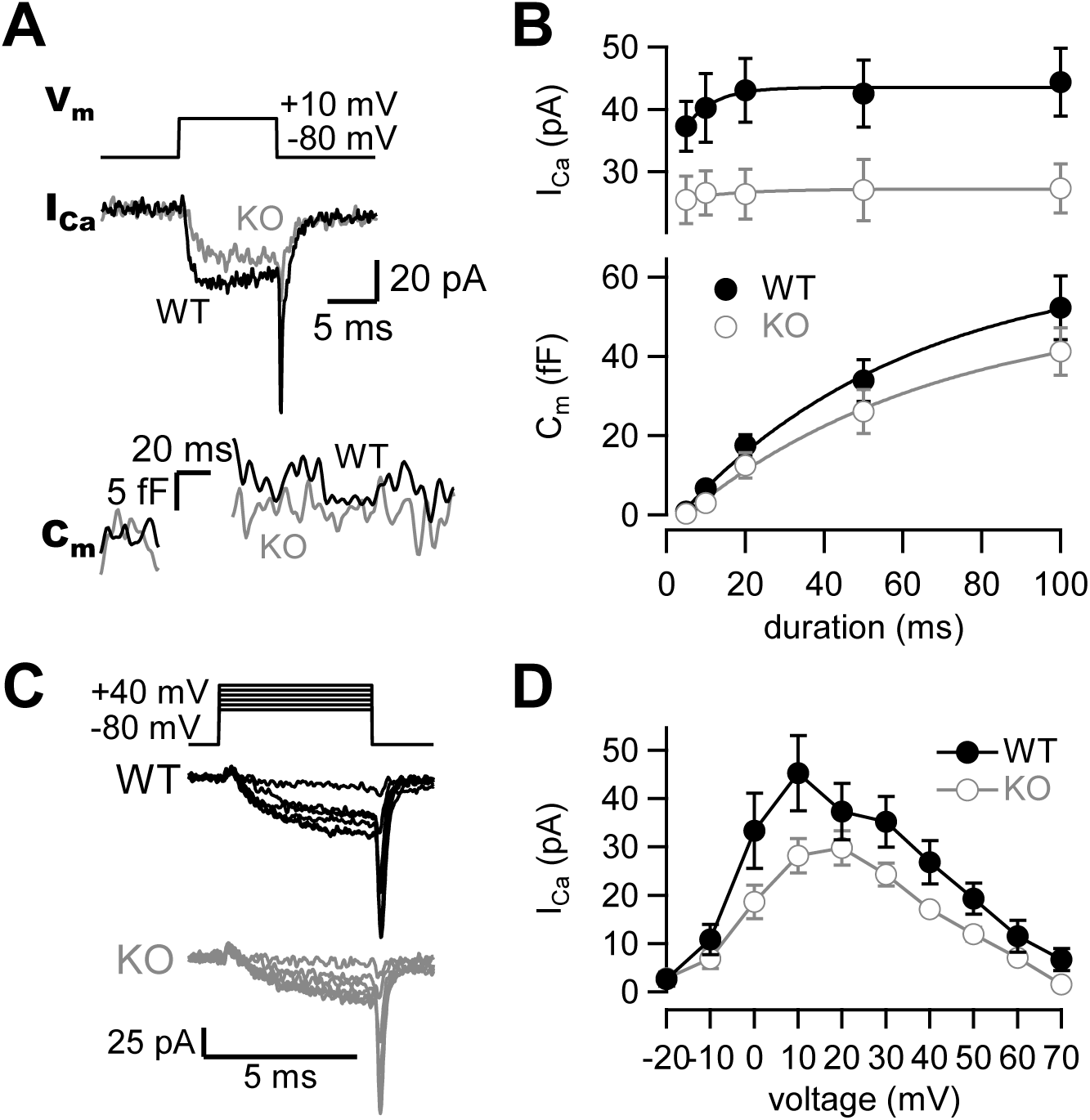
Presynaptic calcium currents and synaptic vesicle release in RIM-BP2 KO mice. (**A**) The terminal was depolarized from -80 mV to +10 mV in WT (black) and RIM-BP2 KO (gray) hippocampal mossy fiber boutons. Ca^2+^ currents (I_Ca_) and membrane capacitance (C_m_) in response to a 10 ms pulse (V_m_) are shown. (**B**) *(top)* The peak Ca^2+^ currents are plotted against the pulse duration. WT vs KO, p = 0.034 (ANOVA). *(bottom)* The capacitance jumps are plotted against the pulse duration. WT vs KO, p = 0.23 (ANOVA). Black filled circles and gray open circles represent the data from WT (n = 9 from 8 animals) and RIM-BP2 KO (n = 8 from 6 animals), respectively. Each data point represents mean ± SEM. (**C**) Experimental protocol and representative traces for presynaptic Ca^2+^ current measurements in WT (black) and RIM-BP2 KO (gray) boutons. Terminals were sequentially depolarized for 5 ms with 2 ms intervals from -80 mV to +70 mV by 10 mV steps. (**D**) Current-voltage relationships of peak Ca^2+^ currents in WT (black filled circle; n = 4-6 from 4 animals) and RIM-BP2 KO (gray open circle; n = 4-5 from 5 animals). I_Ca_s were elicited by 5 ms depolarizations. Two curves were significantly different (p < 0.01 from mixed model). Each data point represents mean ± SEM. Numerical values of plots are provided in Figure 1-source data 1.

**Figure 1-figure supplement 1.**
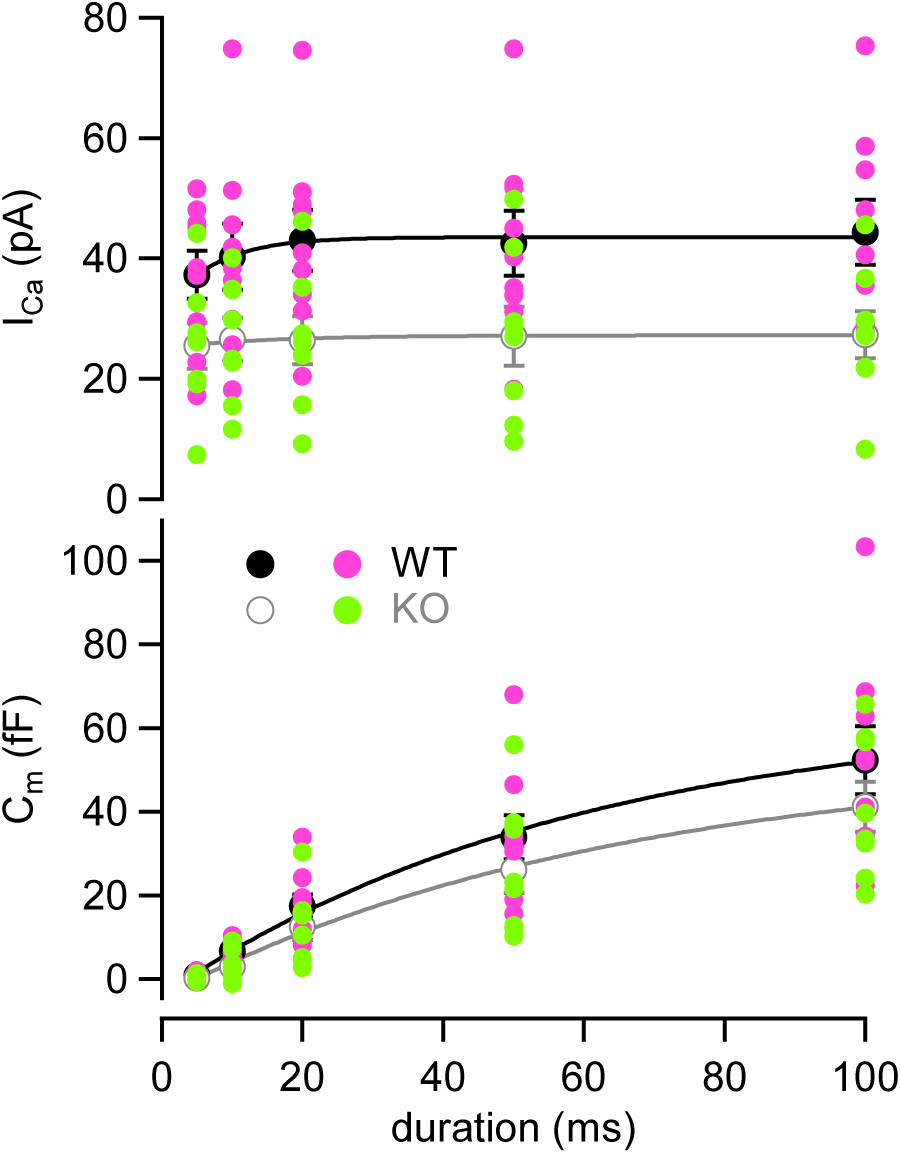
Individual value plot of **Fig. 1B**. Individual values of I_Ca_s (*top*) and C_m_s (*bottom*) in WT (pink) and RIM-BP2 KO (green) terminals are plotted against the pulse duration. Numerical values of plots are provided in Figure 1-source data 1.

**Figure 1-source data 1.** Datasets of Ca^2+^-current and capacitance amplitudes presented in Figure 1 and Figure 1-figure supplement 1.

### The effects of extracellular calcium concentration on Ca^2+^ influx and capacitance changes in WT and RIM-BP2 KO

Because Ca^2+^ currents were reduced in RIM-BP2 KO mice, we tested whether the reduction of Ca^2+^ currents could be restored by elevation of the external Ca^2+^ concentration. We raised the extracellular Ca^2+^ concentration ([Ca^2+^]_ext_) from 2 mM to 4 mM (Fig. 2). Ca^2+^ currents and membrane capacitances were recorded at 4 mM [Ca^2+^]_ext_, and amplitudes then plotted against pulse duration (Fig. 2A and 2-figure supplement 1). Although Ca^2+^ current elicited by a 100 ms pulse in RIM-BP2 KO was still smaller (33 ± 2.7 pA, n = 3) than that in WT (54 ± 6.1 pA, n = 6) (Fig. 2B, top) in 4 mM Ca^2+^, Ca^2+^ currents of KO mice at 4 mM [Ca^2+^]_ext_ were more comparable to those of WT at 2 mM [Ca^2+^]_ext_.

At 4 mM [Ca^2+^]_ext_, ΔC_m_ in response to a 100 ms pulse in RIM-BP2 KO was 51 ± 11 fF (n = 3) and showed a similar value in WT at 2 mM [Ca^2+^]_ext_ (Fig. 2B, bottom). At the same time, ΔC_m_ in WT was not altered (52 ± 2.5 fF, n = 6) at higher [Ca^2+^]_ext_. The time course of exocytosis at 4 mM [Ca^2+^]_ext_ in KO was similar to that of WT at 2 mM [Ca^2+^]_ext_, as the time courses were superimposable in Fig. 2A (p = 0.57 from mixed model).

**Figure 2.**
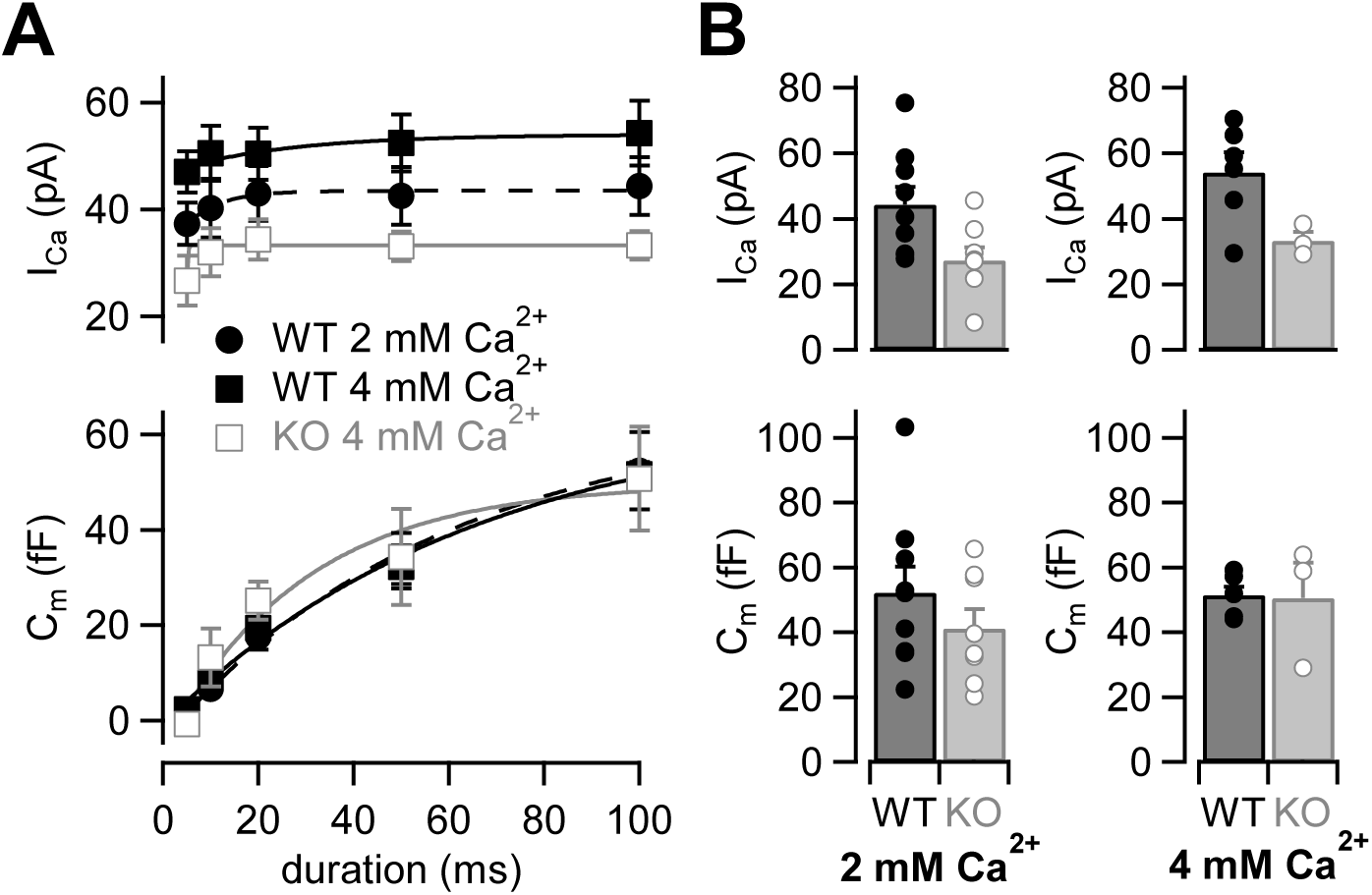
The effects of high extracellular calcium concentration on Ca^2+^ currents and capacitance changes. **(A)** The I_Ca_s (*top*) and the C_m_s (*bottom*) are plotted against the pulse duration. Black filled squares and gray open squares represent the data from WT (n = 6-8 from 6 animals) and RIM-BP2 KO (n = 3-4 from 2 animals) at 4 mM [Ca^2+^]_ext_, respectively. For comparison, the WT data in 2 mM [Ca^2+^]_ext_ are superimposed (black filled circle; n = 9) (the same data sets as Fig. 1B). Each data point represents mean ± SEM. I_Ca_s were significantly different between WT and KO in 4 mM [Ca^2+^]_ext_ (p < 0.01 from mixed model). The WT I_Ca_s in 2 mM [Ca^2+^]_ext_ and the KO I_Ca_s in 4 mM [Ca^2+^]_ext_ were significantly different (p < 0.02 from mixed model), but Cm were not significantly different (p = 0.57 from mixed model). **(B)** Average I_Ca_s *(top)* and C_m_s *(bottom)* in response to a 100 ms pulse in WT (black bars) and RIM-BP2 KO (gray bars) terminals. Extracellular Ca^2+^ concentration was 2 mM *(left)* or 4 mM *(right)*. Error bars show SEM. Circles indicate individual values. Numerical values of plots are provided in Figure 2-source data 1.

**Figure 2-figure supplement 1.**
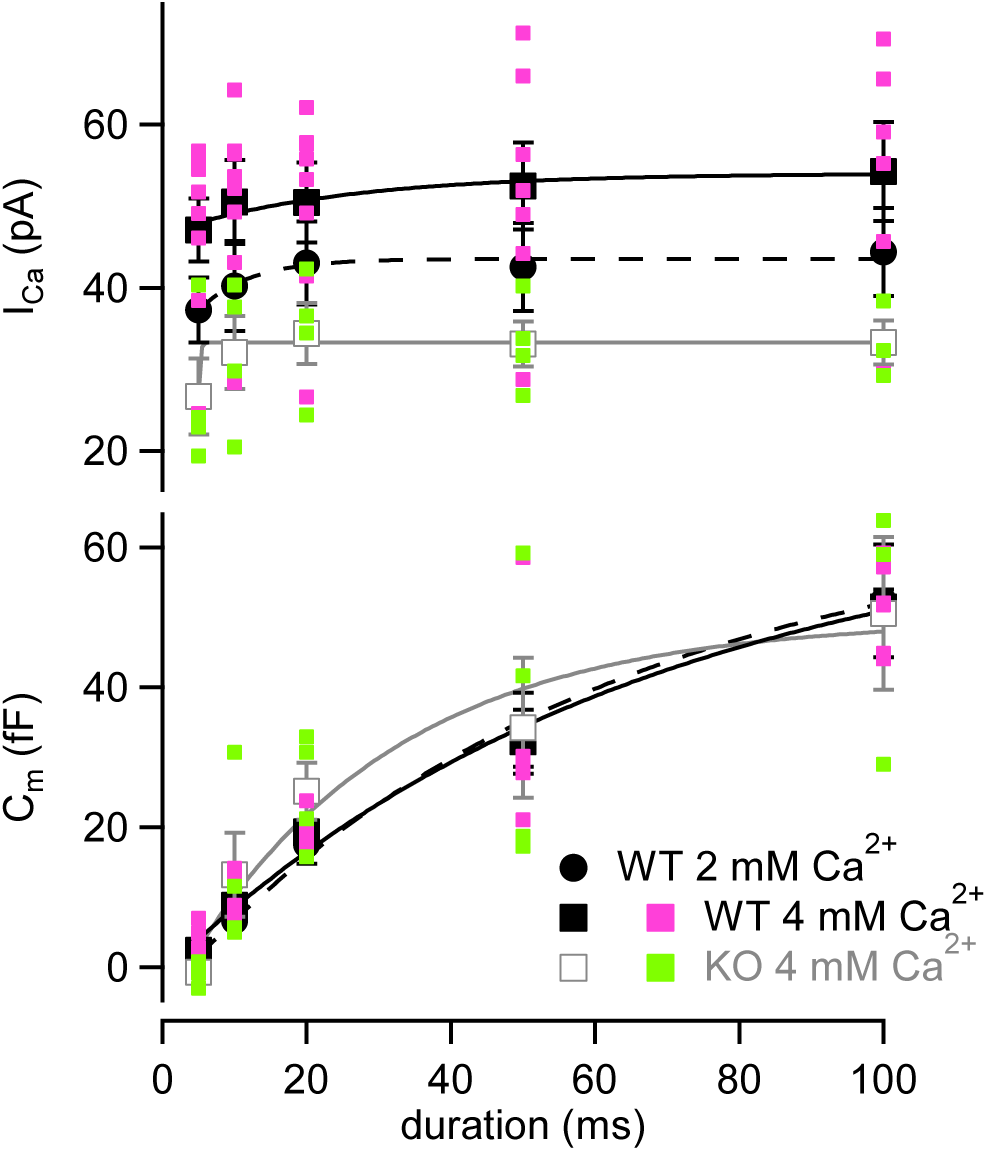
Individual value plot of **Fig. 2A**. Individual values of I_Ca_s (*top*) and C_m_s (*bottom*) recorded at 4 mM [Ca^2+^]_ext_ in WT (pink) and RIM-BP2 KO (green) terminals are plotted against the pulse duration. Numerical values of plots are provided in Figure 2-source data 1.

**Figure 2-source data 1.** Datasets of Ca^2+^-current and capacitance amplitudes presented in Figure 2 and Figure 2-figure supplement 1.

Next, we adjusted Ca^2+^ current amplitudes of WT to that of RIM-BP2 KO by lowering [Ca^2+^]_ext_ (Fig. 3). The amplitudes of Ca^2+^ currents and capacitance jumps were plotted against pulse duration (Fig. 3A and Figure 3-figure supplement 1). At 1 mM [Ca^2+^]_ext_, Ca^2+^ currents and ΔC_m_ were smaller than those of RIM-BP2 KO at 2 mM [Ca^2+^]_ext_. At 1.5 mM [Ca^2+^]_ext_, Ca^2+^ currents became more comparable to those of RIM-BP2 KO at 2 mM [Ca^2+^]_ext_ (p = 0.12 from mixed model). Here, the ΔC_m_ evoked by a 100 ms pulse (43 ± 6.7 fF, n = 5) was comparable between WT and RIM-BP2 KO (Fig. 3A). The average time course of capacitance increase of WT at 1.5 mM [Ca^2+^]_ext_ was similar to that of RIM-BP2 KO at 2 mM [Ca^2+^]_ext_ (p = 0.88 from mixed model). Therefore, by reducing Ca^2+^ currents, WT data could reproduce similar release time course of RIM-BP2 KO. In Fig. 3B, ΔC_m_ in response to a 100 ms pulse at various [Ca^2+^]_ext_s were plotted against the peak Ca^2+^ current amplitudes. The relationship could be fitted by a Hill plot with n = 3 in WT (Schneggenburger et al., 1999). Consistently, when capacitance jumps elicited by various pulse durations are plotted against total Ca^2+^ influx (Figure 3-figure supplement 2), the RIM-BP2 KO data were superimposed on the WT data. In Fig. 3C, relative amplitudes of Ca^2+^ currents and capacitance in response to a 100 ms pulse are plotted. The RIM-BP2 KO data were superimposed on the WT data, suggesting that the RRP size were similar between WT and KO (∼50 fF). Taken together, presynaptic capacitance measurements were not able to detect differences in the release rates between WT and KO especially when Ca^2+^ influx in WT and KO was adjusted to be similar. In particular, the RRP sizes measured by the pool depleting stimulation were not altered strongly by RIM-BP2 KO. The results may mean that RIM-BP2 is dispensable for transmitter release or that large Ca^2+^, or large Ca^2+^ influx may overcome the deficit of the release. Alternatively, capacitance measurements may not be sensitive enough to detect the differences between WT and KO at short stimulation, which are relevant for AP-evoked release (see below, Fig. 5 and Fig. 6).

**Figure 3.**
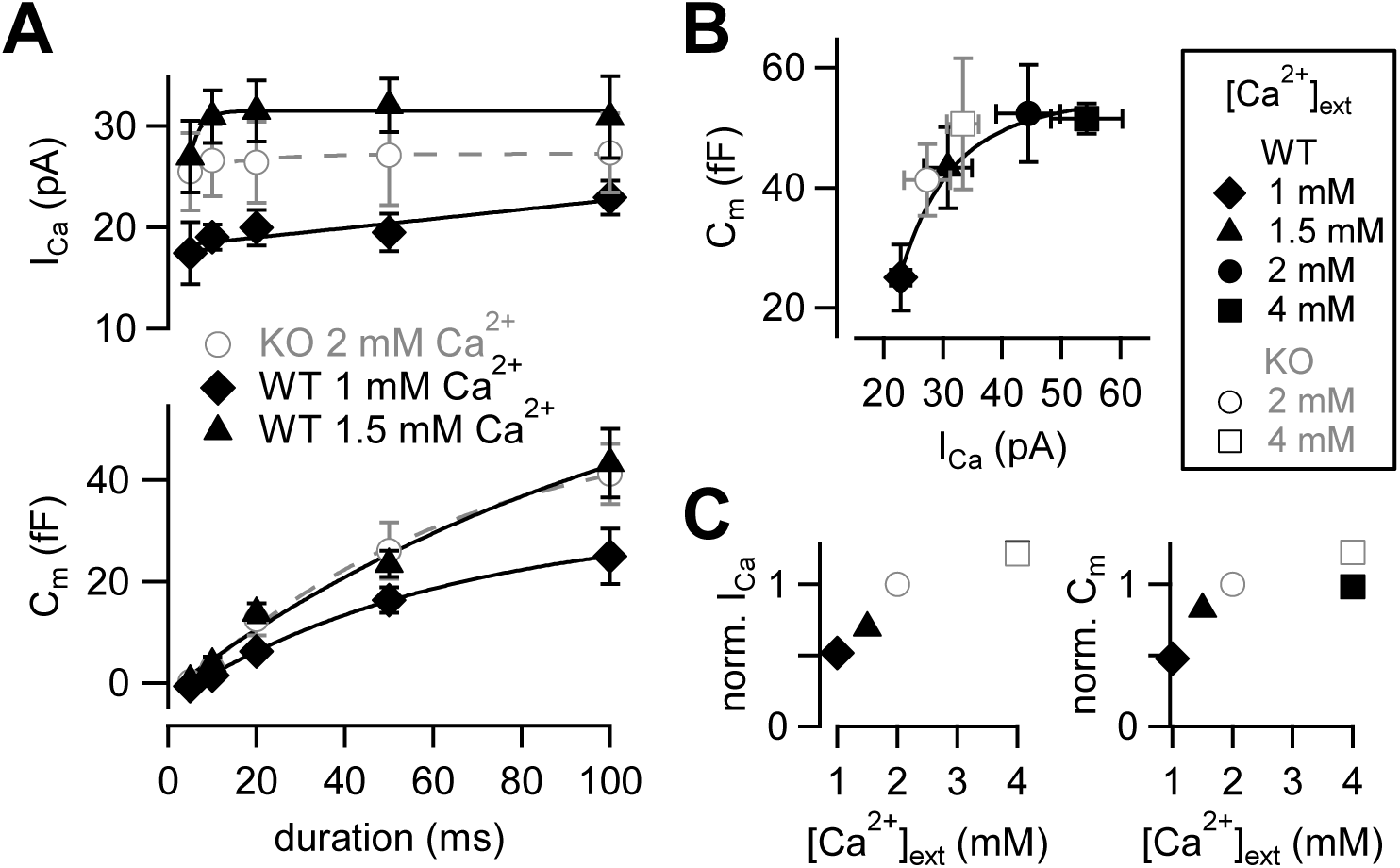
Calcium-dependence of the release kinetics and the RRP size. (**A**) *(top)* The relationship between peak Ca^2+^ currents and pulse durations at different [Ca^2+^]_ext_s. *(bottom)* The relationship between capacitance jumps and pulse durations at different [Ca^2+^]_ext_s. Gray open circles, black diamonds and black triangles represent the data from RIM-BP2 KO at 2 mM [Ca^2+^]_ext_ (n = 8) (the same data sets as Fig. 1B), WT at 1 mM [Ca^2+^]_ext_ (n = 3-5 from 4 animals) and WT at 1.5 mM [Ca^2+^]_ext_ (n = 5-7 from 6 animals), respectively. Each data point represents mean ± SEM. (**B**) Capacitance jumps at various [Ca^2+^]_ext_s are plotted against calcium current amplitudes. Pulses were 100 ms depolarization from -80 mV to +10 mV. Each data point represents mean ± SEM. Data points were fitted with a Hill equation with n = 3. (**C**) *(left)* Average I_Ca_s at indicated [Ca^2+^]_ext_s were normalized to the response at 2 mM [Ca^2+^]_ext_ in each genotype. *(right)* Average C_m_s at indicated [Ca^2+^]_ext_s were normalized to the 2 mM [Ca^2+^]_ext_ response in each genotype. Numerical values of plots are provided in Figure 3-source data 1.

**Figure 3-figure supplement 1.**
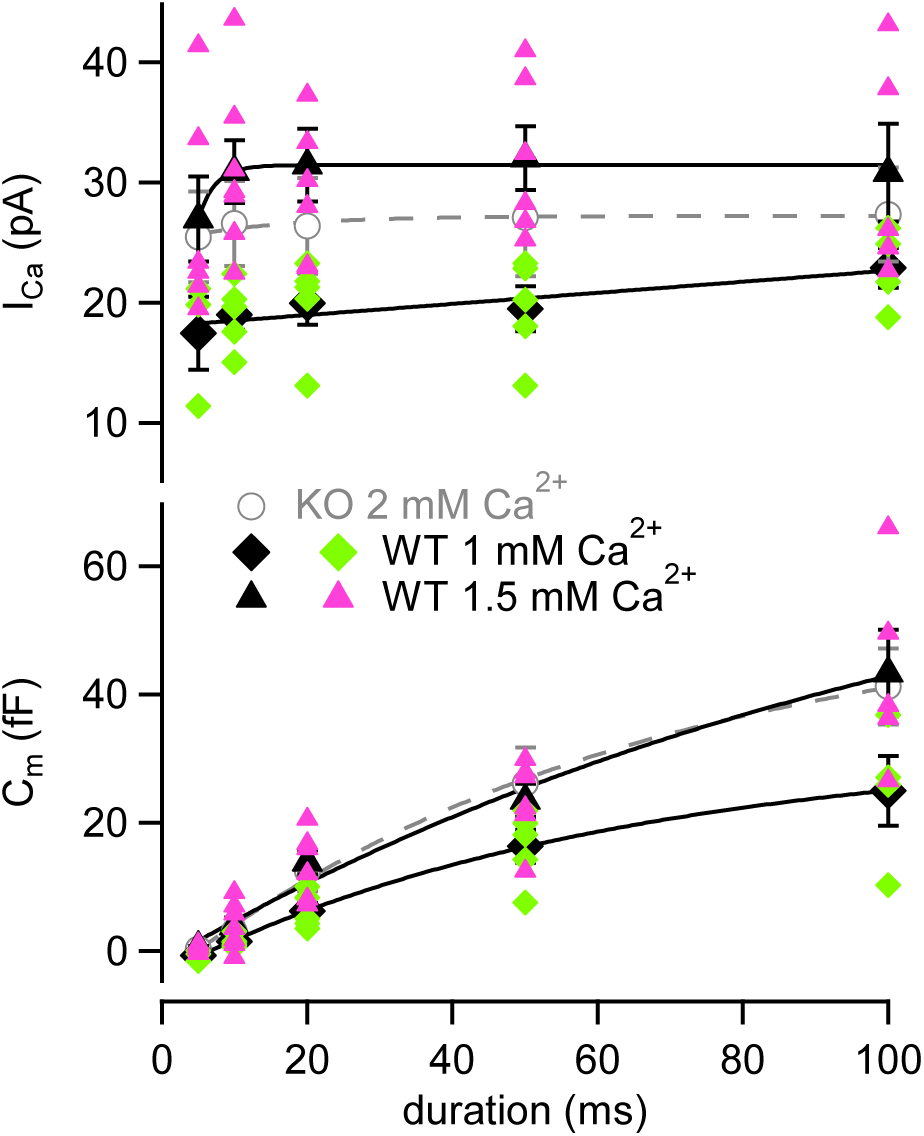
Individual value plot of **Fig. 3A**. Individual values of I_Ca_s (*top*) and C_m_s (*bottom*) recorded at 1 mM (green) or 1.5 mM (pink) [Ca^2+^]_ext_ in WT are plotted against the pulse duration. Numerical values of plots are provided in Figure 3-source data 1.

**Figure 3-source data 1.** Datasets of numerical values presented in Figure 3 and Figure 3-figure supplement 1,

**Figure 3-figure supplement 2.**
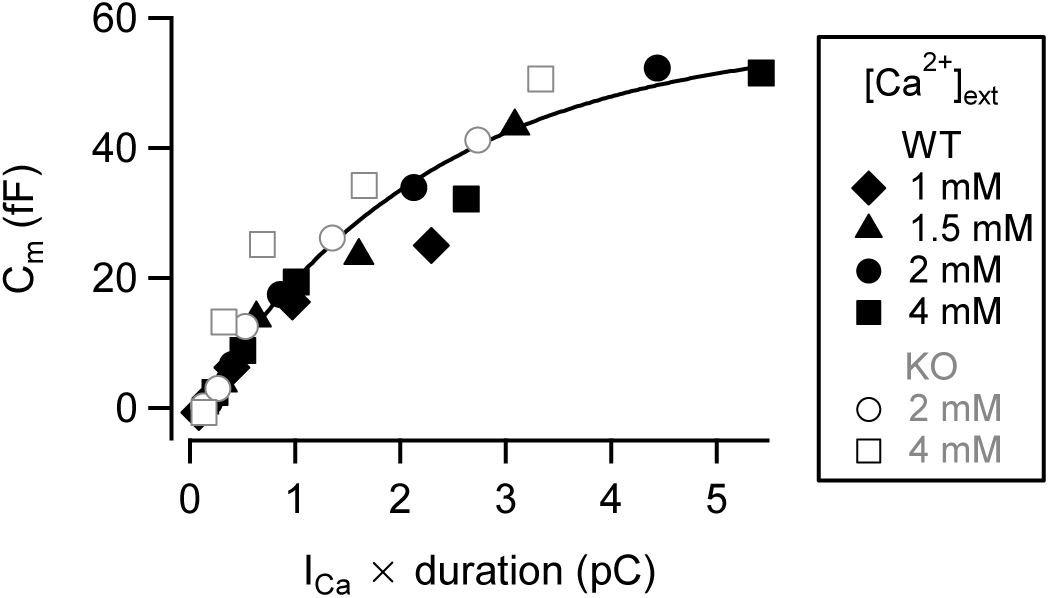
The relationship between synaptic vesicle release and total Ca^2+^ charge. Capacitance jumps elicited by various pulse durations are plotted against total Ca^2+^ charge. The total Ca^2+^ charge was calculated by multiplying a peak Ca^2+^ current amplitude by pulse duration. Each symbol represents a data point obtained from WT (black filled symbols) and RIM-BP2 KO (gray open symbols) mice at various [Ca^2+^]_ext_. Data points were fitted by a single exponential function. Numerical values of plots are provided in Figure 3-figure supplement 2-source data 1.

**Figure 3-figure supplement 2-source data 1.** Datasets of Ca^2+^ charge and ΔC_m_ presented in Figure 3-figure supplement 2.

### High EGTA experiments suggest unaltered coupling between calcium channels and synaptic vesicles

Previous studies have used Ca^2+^ chelator such as EGTA to examine the sensitivity of release to intracellular Ca^2+^ buffers (Adler et al., 1991; Borst & Sakmann, 1996; Vyleta & Jonas, 2014). When the physical coupling between Ca^2+^ channels and synaptic vesicles is tight, EGTA (slow Ca^2+^ chelator) has less effect on release probability, but EGTA is effective when the coupling is loose. Previous experiments have suggested that RIM-BPs regulated the channel-vesicle coupling at the calyx of Held (Acuna et al., 2015). If this observation also applies to the mossy fiber synapse, EGTA sensitivity of release should be higher in RIM-BP2 KO terminals. To investigate the coupling at mossy fiber boutons in RIM-BP2 KO, we changed the concentration of EGTA in the patch pipette. In the presence of 5 mM EGTA, we depolarized the terminal from -80 mV to +10 mV for 5 ms and recorded Ca^2+^ currents and membrane capacitance (Fig. 4A). Ca^2+^ currents and ΔC_m_ were plotted against pulse duration (Fig. 4B and Figure 4-figure supplement 1). In both WT and RIM-BP2 KO mice, ΔC_m_ were reduced under 5 mM EGTA compared with the control condition of 0.5 mM EGTA. We compared the effect of EGTA on average amplitudes of peak Ca^2+^ currents in response to a 20 ms pulse (Fig. 4D). Interestingly, Ca^2+^ current amplitudes were ∼1.2 times larger at 5 mM EGTA (53 ± 12 pA, n = 5) than at 0.5 mM EGTA (43 ± 5.1 pA, n = 9) in WT, though the data scattered under 5 mM EGTA condition and there was no statistically significant difference (p = 0.36, t-test). In RIM-BP2 KO, we did not observe such an increase. EGTA might inhibit Ca^2+^ channel inactivation in WT (von Gersdorff and Matthews, 1996). In Fig. 4C, we compared the effect of EGTA on the ΔC_m_ in response to a 20 ms pulse. In both genotypes, ΔC_m_ values were inhibited by 5 mM EGTA, and the inhibition was somewhat larger in RIM-BP2 KO (EGTA inhibition: p < 0.01, genotypes p > 0.05 from mixed model).

Ca^2+^ current amplitudes in RIM-B2 KO are smaller than in WT (Fig. 4B). It is possible that strong effect of high EGTA on release may be due to reduced Ca^2+^ currents in RIM-BP2 KO. Therefore, we changed [Ca^2+^]_ext_ from 2 mM to 1 mM and adjusted Ca^2+^ current amplitudes of WT to the RIM-BP2 KO level. The capacitance increase and the amplitudes of Ca^2+^ currents became similar to those of RIM-BP2 KO (Fig. 4C and Fig. 4D). These results indicated that WT and RIM-BP2 KO terminals had similar sensitivity of neurotransmitter release to EGTA. Thus, it is unlikely that changes in the coupling distance are responsible for the reduced exocytosis of the mutants.

**Figure 4.**
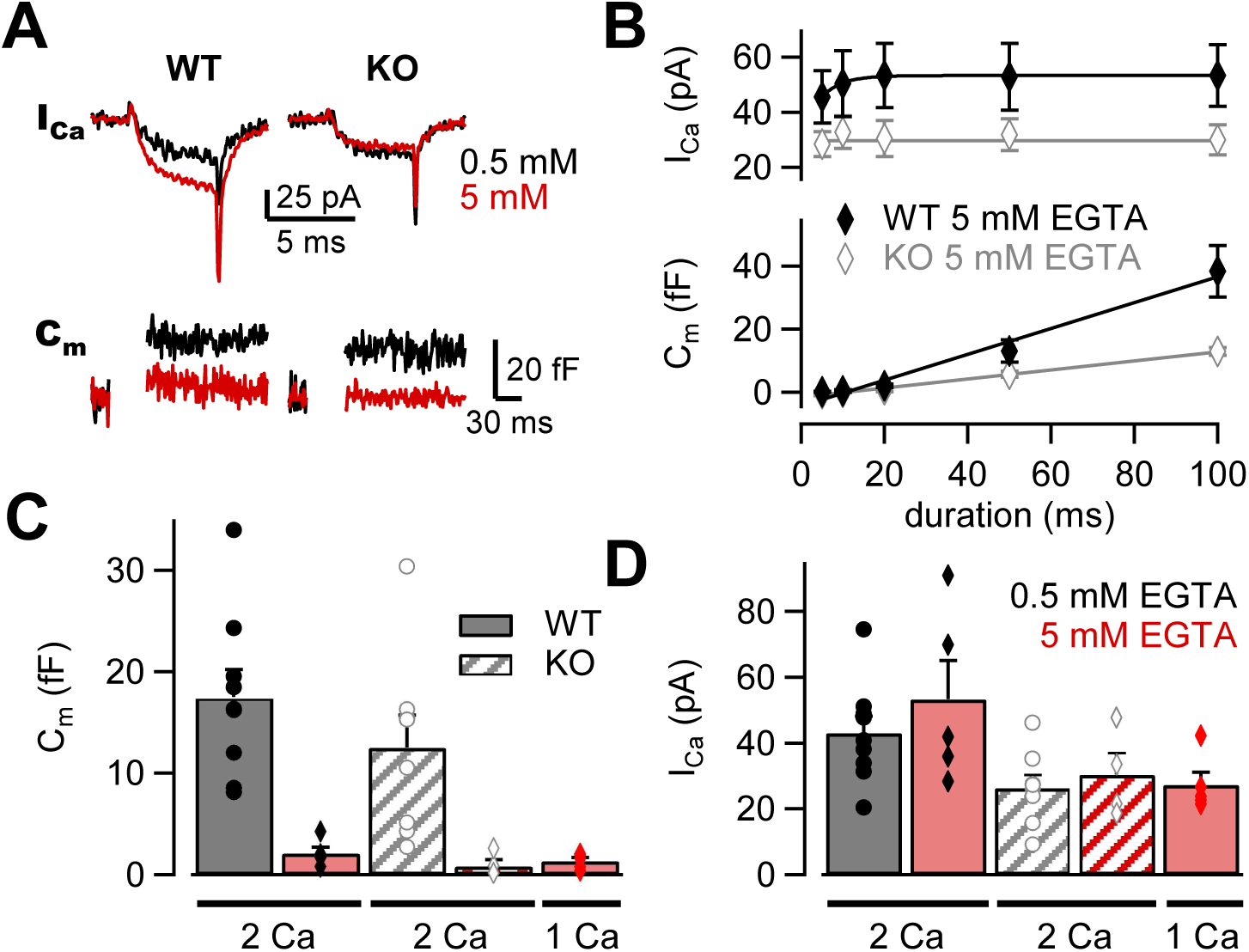
The effects of high EGTA on calcium currents and synaptic vesicle exocytosis. (**A**) Example traces in response to a 5 ms depolarizing pulse to +10 mV in WT *(left)* and RIM-BP2 KO *(right)* boutons. Ca^2+^ current (I_Ca_) and membrane capacitance (C_m_) recorded with 0.5 mM EGTA (black) or 5 mM EGTA (red) in the patch pipette are shown. Note that each trace was obtained from different terminals but the traces were superimposed. (**B**) The I_Ca_s *(top)* and C_m_s *(bottom)* are plotted against the pulse duration. The patch pipette contained 5 mM EGTA. Black filled diamonds and gray open diamonds represent the data from WT (n = 4-5 from 4 animals) and RIM-BP2 KO (n = 4 from 3 animals), respectively. Each data point indicates mean ± SEM. (**C, D**) Average C_m_s (**C**) and I_Ca_s (**D**) elicited by a 20 ms pulse in WT (filled bars) and RIM-BP2 KO (hatched bars). The extracellular Ca^2+^ concentration was 1 mM (from 5 animals) or 2 mM. The intracellular solution contained either 0.5 mM EGTA (black) or 5 mM EGTA (red). Error bars show SEM. Circles and diamonds indicate individual values. Numerical values of plots are provided in Figure 4-source data 1.

**Figure 4-figure supplement 1.**
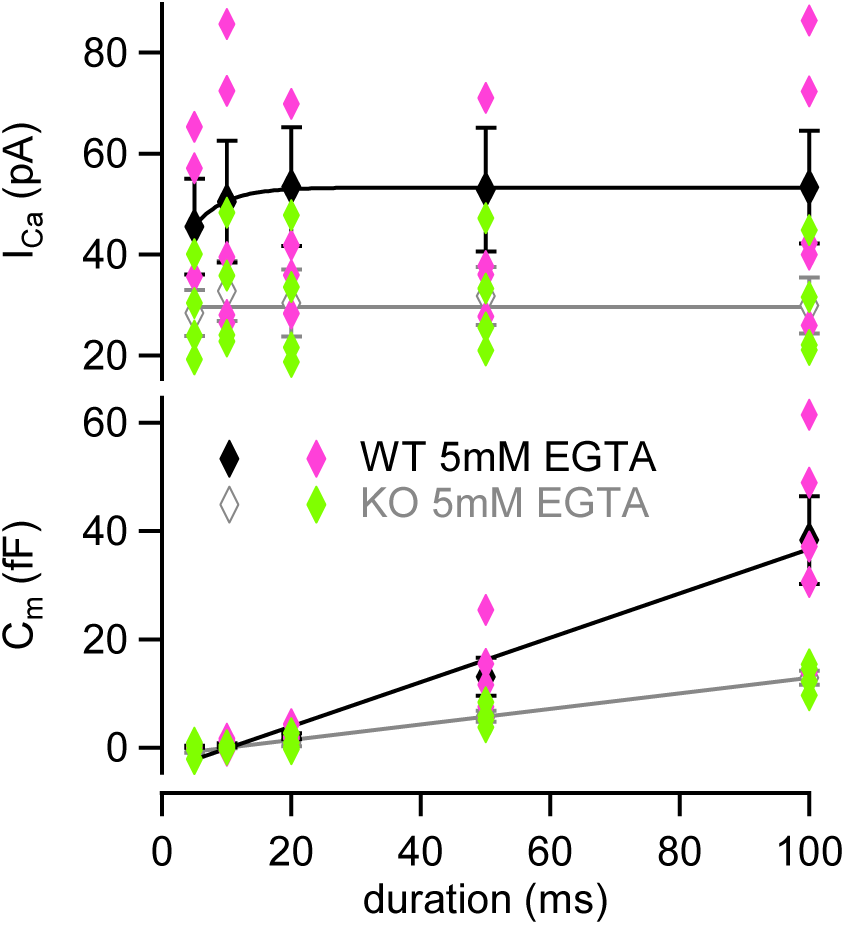
Individual value plot of Fig. 4B. Individual values of I_Ca_s (*top*) and C_m_s (*bottom*) recorded in the presence of 5 mM EGTA in WT (green) and KO (pink) were plotted against the pulse duration. Numerical values of plots are provided in Figure 4-source data 1.

**Figure 4-source data 1.** Datasets of Ca^2+^-current and capacitance amplitudes presented in Figure 4 and Figure 4-figure supplement 1.

### Smaller AP-evoked EPSCs in KO could not be explained entirely by the reduced Ca^2+^ influx

While Ca^2+^ current amplitudes were clearly reduced in KO, capacitance jumps were reduced little. Capacitance measurements could only detect large amounts of exocytosis (∼5 fF, 50 vesicles), and the changes relevant for the AP-evoked release (< 10 vesicles per AP per synapse) could not be detected. Moreover, large Ca^2+^ influx used for the stimulation protocol of Fig.1 to Fig. 3 could overcome the deficits of RIM-BP2 KO. In order to detect the changes related to physiological conditions, we measured the AMPA-EPSCs evoked by electrical mossy fiber stimulation at CA3 pyramidal cells (Fig. 5). As seen in Fig. 5A and Fig. 5B, the EPSCs were evoked by dual stimulation of mossy fibers. The mossy-fiber evoked responses were confirmed by the relatively faster EPSC kinetics and sensitivity to DCG-Ⅳ (1 μM), a mGluR2 agonist (remaining response of < 30 %, 0.23 ± 0.01 for WT (n = 5), 0.29 ± 0.02 for KO (n = 5)). Because multiple fibers were stimulated, we could not estimate the number of vesicles released per one synapse. Nevertheless, when fibers were stimulated similarly, the EPSC amplitudes of KO were much smaller (893 ± 202 pA in WT, n = 7; 189 ± 73 pA in KO, n = 7) (p = 0.011). In Fig. 2, Presynaptic recordings indicated that Ca^2+^ current amplitudes of KO at 4 mM [Ca^2+^]_ext_ became similar to those of WT at 2 mM [Ca^2+^]_ext_ (Fig. 2). The EPSCs became larger when the [Ca^2+^]_ext_ was raised from 2 mM to 4 mM in KO (Fig. 5A, top, see also Figure 6-figure supplement 1) and became more similar to those of WT. In Fig. 3, presynaptic recording showed that I_Ca_s of WT at 1.5 mM [Ca^2+^]_ext_ were similar to those of KO at 2 mM [Ca^2+^]_ext_. However, the WT EPSCs at 1.5 mM [Ca^2+^]_ext_ seemed larger than KO at 2 mM [Ca^2+^]_ext_ (Fig. 5A, bottom, see also Figure 6-figure supplement 1). Changing the [Ca^2+^]_ext_ from 1.5 mM to 2 mM in WT and from 2 mM to 4 mM in KO should have the same potentiation effect on the EPSCs, if the reduced Ca^2+^ current amplitudes are the only mechanism for reduced EPSCs in KO. In Fig. 5B, we calculated two types of potentiation ratios: In WT, the ratio of the EPSC response in 2 mM to that in 1.5 mM [Ca^2+^]_ext_ was calculated in each cell. In KO, the ratio of the EPSC response in 4 mM to that in 2 mM [Ca^2+^]_ext_ was calculated in each cell. When these two values were compared, the ratios were much larger in KO (Fig. 5B, p < 0.01, t-test). Pr is more sensitive to the [Ca^2+^]_ext_ in the low end of the dose-response curve because of the 3^rd^-4^th^ power dependence of release on [Ca^2+^]_ext_ at the low end (Dodge and Rahamimoff, 1967; Schneggenburger and Neher, 2000; Bollmann et al., 2000). Therefore, the result suggests lower release probability in KO, even when the amounts of Ca^2+^ influx were adjusted to be similar. When the paired pulse ratios were compared, all four conditions (WT in 1.5 mM and 2 mM [Ca^2+^]_ext_, KO in 2 mM and 4 mM [Ca^2+^]_ext_) provided similar values (Fig. 5C). The paired pulse ratio should be inversely correlated to Pr, in principle. However, paired pulse facilitation in mossy fiber synapses could be mediated by multiple effects, including Ca^2+^-independent broadening of AP waveforms (Geiger and Jonas, 2000) and Ca^2+^ buffer saturation which is more pronounced and increases the paired-pulse ratios at higher [Ca^2+^]_ext_ (Blatow et al., 2003). Therefore, the value might not be sensitive to the [Ca^2+^]_ext_.

We also analyzed the EPSCs in response to a stimulus train (50 Hz × 25 or 26 pulses) both in WT and KO (Fig. 6) under different external Ca^2+^ concentrations. When the WT and KO responses were compared, the time course of facilitation/depression was quite different between WT and KO (Fig. 6A-Fig. 6C). In 2 mM [Ca^2+^]_ext_, WT responses showed initial facilitation (∼2 fold) followed by depression, whereas KO responses showed pronounced facilitation of ∼10 fold. The difference could be due to both reduced Ca^2+^ currents and impaired fusion competence in KO.

**Figure 5.**
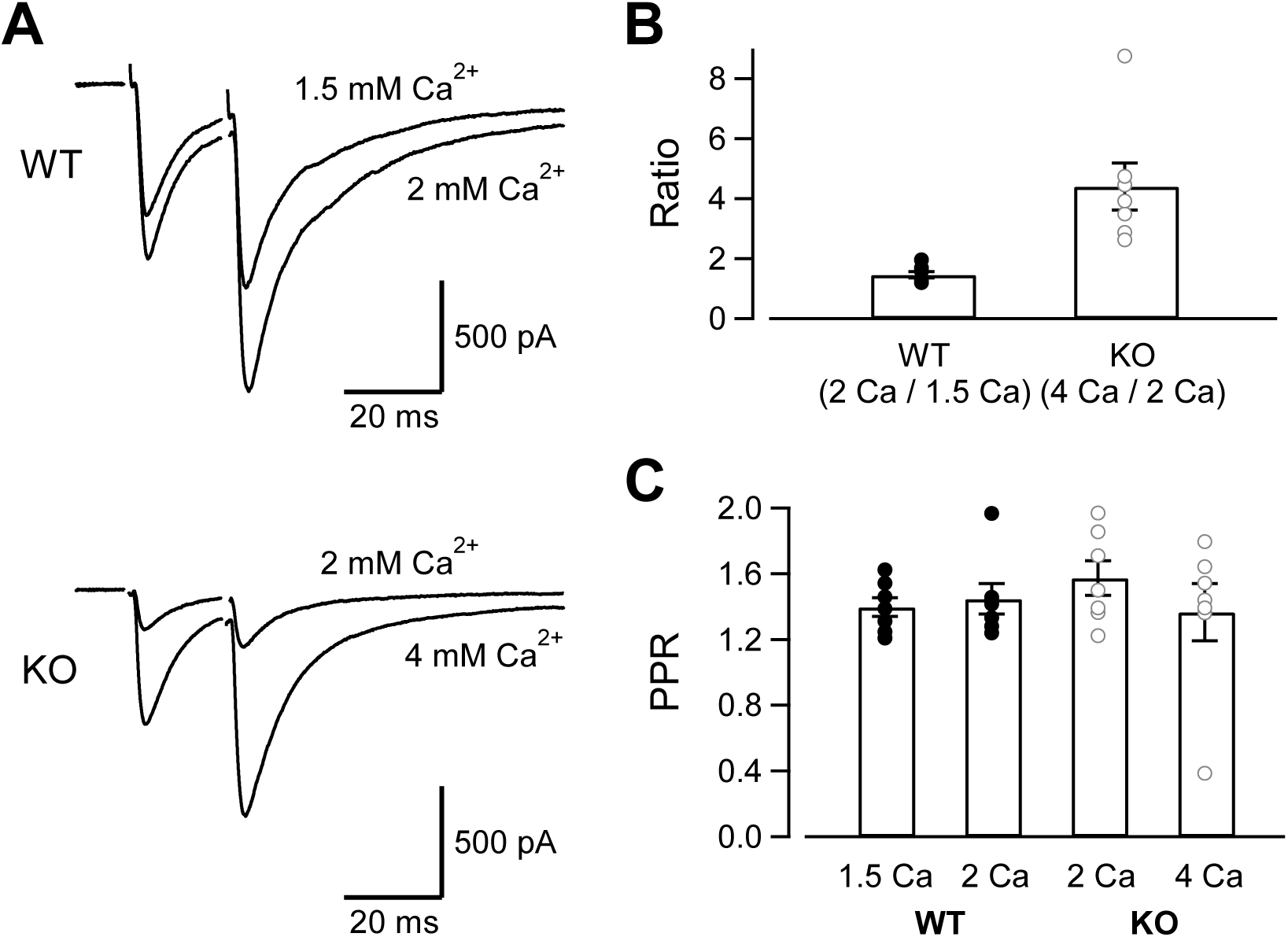
The evoked EPSC amplitudes in WT and KO, and their sensitivity to the extracellular Ca^2+^. **(A)** The mossy fiber-evoked EPSCs measured at CA3 pyramidal cells. The fibers were stimulated twice with an interval of 20 ms. The responses of WT in 1.5 mM and 2 mM [Ca^2+^]_ext_ (n = 7 cells from 5 animals), as well as those of KO in 2 mM and 4 mM [Ca^2+^]_ext_ (n = 7 cells from 6 animals) are shown. The stimulus artifacts were blanked. **(B)** The amplitude ratios of WT EPSCs (2 mM / 1.5 mM [Ca^2+^]_ext_) and KO EPSCs (4 mM / 2 mM [Ca^2+^]_ext_) are shown. The concentrations were chosen to set the Ca^2+^ influx similar between WT and KO from Fig. 1-Fig. 3. **(C)** The paired pulse ratios under 4 conditions (WT in 1.5 mM and 2 mM [Ca^2+^]_ext_, KO in 2 mM and 4 mM [Ca^2+^]_ext_). Numerical values of plots are provided in Figure 5-source data 1.

**Figure 5-source data 1.** Datasets of numerical values presented in Figure 5.

To adjust presynaptic Ca^2+^ influx to the similar level as the KO terminals, the [Ca^2+^]_ext_ was reduced to 1.5 mM in WT. Under this condition, presynaptic Ca^2+^ current amplitudes were expected to be similar between WT and KO (in 2 mM [Ca^2+^]_ext_). If the reduced Ca^2+^ influx is only responsible for smaller EPSCs in KO, we will observe similar time course of train responses between WT and KO. However, facilitation was much less in the WT condition (at 1.5 mM [Ca^2+^]_ext_, Fig. 6D, p < 0.05 from ANOVA), suggesting the mechanism independent of Ca^2+^ influx. In order to obtain the same Ca^2+^ influx as that of WT in 2 mM [Ca^2+^]_ext_, the [Ca^2+^]_ext_ was increased to 4 mM in KO. The time course of facilitation was somewhat larger in the KO condition compared with that WT in 2 mM [Ca^2+^]_ext_. Because of the data scatter in KO, the time course was statistically not significant (p > 0.05 from ANOVA). Because multiple fibers were stimulated, the absolute EPSC amplitudes depend on not only the release probability and the number of release sites per synapse, but also the number of stimulated synapses. Nevertheless, when the absolute amplitudes were plotted against the stimulus number, the differences were most prominent at the first responses in the train and the differences were diminished later in the train, suggesting that the initial release probability was likely to be lower at RIM-BP2 KO synapses, but the release could be restored during repetitive stimulation (Figure 6-figure supplement 1). Restoration of release is consistent with minor difference of capacitance changes in response to the depolarization (Figs. 1-4).

**Figure 6.**
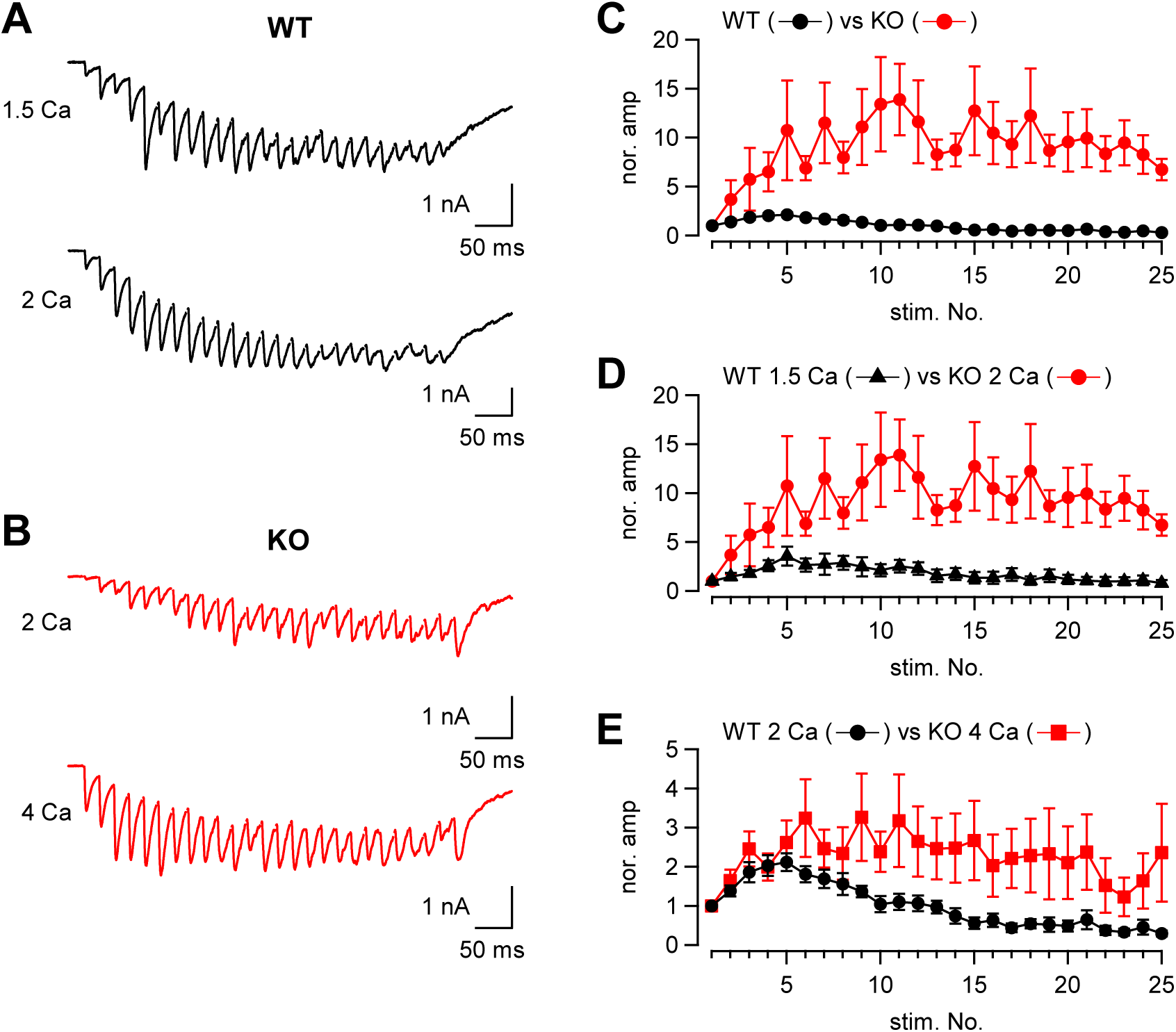
The time course of synaptic facilitation/depression in WT and KO. **(A)** Mossy fibers were stimulated at 50 Hz (25 times) and the evoked EPSCs were measured in 1.5 mM (n = 5 cells from 4 animals) and 2 mM [Ca^2+^]_ext_ (n = 7 cells from 5 animals) in WT. The stimulus artifacts were blanked. **(B)** Mossy fibers were stimulated at 50 Hz (26 times in this particular example), and the evoked EPSCs were measured in 2 mM (n = 7 cells from 6 animals) and 4 mM [Ca^2+^]_ext_ (n = 8 cells from 6 animals). **(C)** The time course of EPSC amplitudes during a 50 Hz train. The data were obtained from WT and KO in 2 mM [Ca^2+^]_ext_. The time courses were significantly different (p < 0.01 from ANOVA). **(D)** The same as **(C)**, but the data were obtained from WT in 1.5 mM [Ca^2+^]_ext_ and KO in 2 mM [Ca^2+^]_ext_. The time courses were significantly different (p < 0.05 from ANOVA). **(E)** The same as **(C)**, but the data were obtained from WT in 2 mM [Ca^2+^]_ext_ and KO in 4 mM [Ca^2+^]_ext_. Numerical values of plots are provided in Figure 6-source data 1. The time courses were not significantly different (p = 0.12 from ANOVA). Numerical values of plots are provided in Figure 6-source data 1.

**Figure 6-figure supplement 1.**
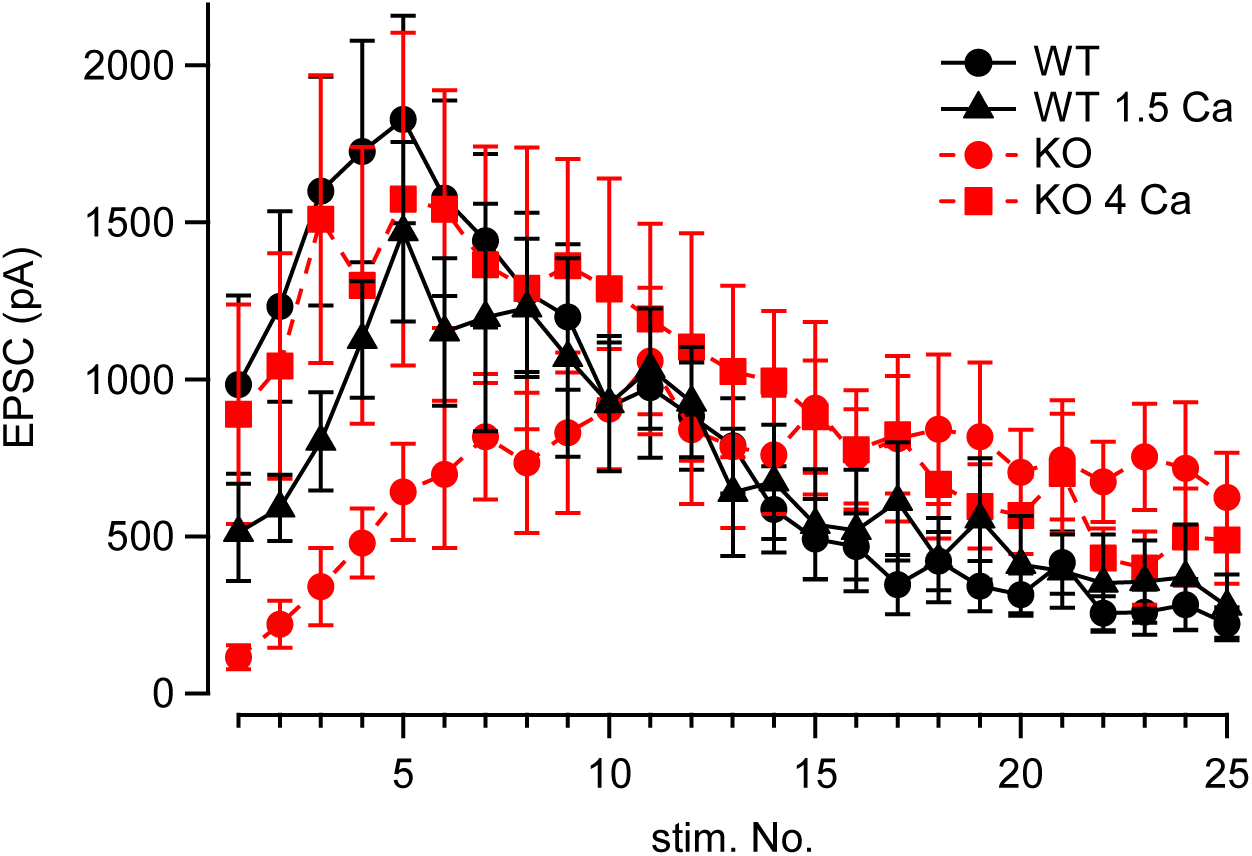
The EPSC amplitudes plotted against the stimulus number. The same data sets as Fig. 6C-Fig. 6E, but the absolute EPSC amplitudes are plotted instead of the normalized amplitudes. Numerical values of plots are provided in Figure 6-source data 1. Raw data were summarized. In Fig. 5 and Fig. 6, the effect of holding potential was corrected so that the values were adjusted to those at -70 mV. Numerical values of plots are provided in Figure 6-source data 1.

**Figure 6-figure supplement 2.**
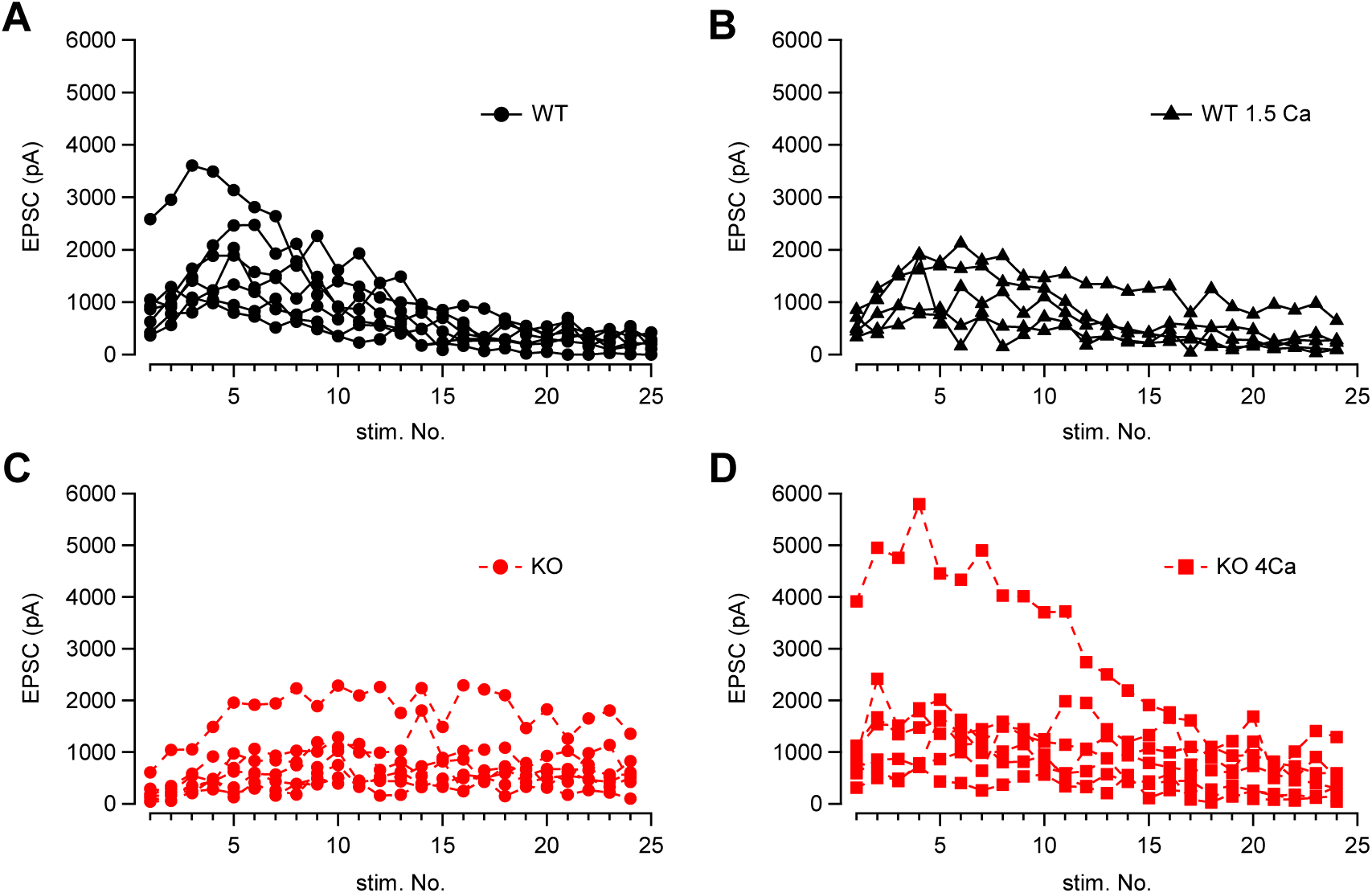
The individual data of **Fig. 6**. WT in 2mM (**A**) and 1.5 mM (**B**) [Ca^2+^]_ext_, and KO in 2 mM (**C**) and 4 mM [Ca^2+^]_ext_ (**D**) are shown. Numerical values of plots are provided in Figure 6-source data 1.

**Figure 6-source data 1.** Datasets of numerical values presented in Figure 6, Figure 6-figure supplement 1, and Figure 6-figure supplement 2.

### STED microscopy is consistent with less abundance of P/Q-type Ca^2+^ channels in RIM-BP2 KO

Our electrophysiological data suggest that RIM-BP2 may regulate the abundance of Ca^2+^ channels. In order to directly study the Ca^2+^ channel abundance at the AZ, we performed STED microscopy analysis of Ca^2+^ channels in WT and RIM-BP2 KO hippocampal mossy fiber terminals. We here focused on P/Q-type and N-type Ca^2+^ channels because both Ca^2+^ channel types are relevant for transmitter release at this synapse (Castillo et al., 1994; Pelkey et al., 2006; Li et al., 2007). We performed immunohistochemistry on thin brain cryosections from WT and RIM-BP2 KO mice. STED microscopy confirmed complete loss of RIM-BP2 proteins in KO terminals (Figure 7-figure supplement 1). We then immunohistochemically labeled the α-subunit of either P/Q-type Ca^2+^ channel (Cav2.1) or N-type Ca^2+^ channel (Cav2.2) and Munc13-1 (AZ/release site marker, Böhme et al., 2016; Sakamoto et al., 2018) with VGLUT1 (presynaptic marker) and PSD-95 (postsynaptic marker) (Fig. 7A, Fig. 7B and Figure 7-figure supplement 2). In the stratum lucidum of the hippocampal CA3 region, mossy fibers form large synapses containing multiple and clustered AZs on CA3 pyramidal cells, and also small en passant and filopodial synapses containing a single AZ on interneurons (Acsády et al., 1998; Rollenhagen et al., 2007). Consistent with this, large VGLUT1 positive terminals rarely contacted interneurons marked with two major interneuron markers, mGluR1α and GluA4 (Figure 7-figure supplement 3). Thus, we identified AZs using Munc13-1 STED images and quantified the signal intensity of Cav2.1 or Cav2.2 only at AZs clustered in large VGLUT1 positive terminals in the CA3 stratum lucidum, allowing us to confine the analysis to large mossy fiber terminals innervating onto CA3 pyramidal cells. The total signal intensity of Cav2.1 at the AZ was 22% lower in RIM-BP2 KO mice (n = 4 animals) than in WT mice (n = 4 animals) (p = 0.0286) (Fig. 7C, left). These data are consistent with the reduction of Ca^2+^ currents in RIM-BP2 KO mice (Fig. 1), though STED microscopy is non-linear and can only provide qualitative differences regarding intensity. In addition, the spatial resolution of STED microscopy (especially z axis) does not assign the signals to the large mossy fiber terminal rigidly. The area, length, and width of Cav2.1 clusters in the AZ were not different between WT and RIM-BP2 KO terminals (Figure 7-figure supplement 4). The analysis in this study might not have spatial resolution to detect the differences. The analysis from the confocal microscopy also indicated that the signal intensity of Cav2.1 was reduced in KO, consistent with the STED data (Figure 7-fugure supplement 5).

In contrast to Cav2.1, the total signal intensity of Cav2.2 and Munc13-1 did not significantly differ between WT and RIM-BP2 KOs (p = 0.3143 and p = 0.0831) (Fig. 7C, right), though it remains possible that we failed to detect some changes. Furthermore, we found that the number of Cav2.1 clusters within the AZ identified by STED deconvolution analysis was not altered (2.5 ± 0.11 in WT, n = 4 animals; 2.3 ± 0.02 in KO, n = 4 animals) (p = 0.114). These data are consistent with the idea that RIM-BP2 KO reduces the abundance of P/Q-type Ca^2+^ channels per cluster. To optically estimate physical distances between Ca^2+^ channels and release sites, we next analyzed the nearest neighboring distance between Cav2.1 and Munc13-1 clusters. These distances were unchanged (61 ± 1.4 nm in WT, n = 4 animals; 64 ± 1.3 nm in KO, n = 4 animals) (p = 0.200) (Fig. 7D), consistent with the similar sensitivity of release to EGTA between WT and RIM-BP2 KO mice. However, the resolution of STED microscopy cannot allow detection of small differences of the cluster number and the distance between WT and KO. From these results together with presynaptic recordings, we suggest that decreased Ca^2+^ influx in RIM-BP2 KO hippocampal mossy fiber terminals is caused by less abundance of P/Q-type Ca^2+^ channels at the AZ.

**Figure 7.**
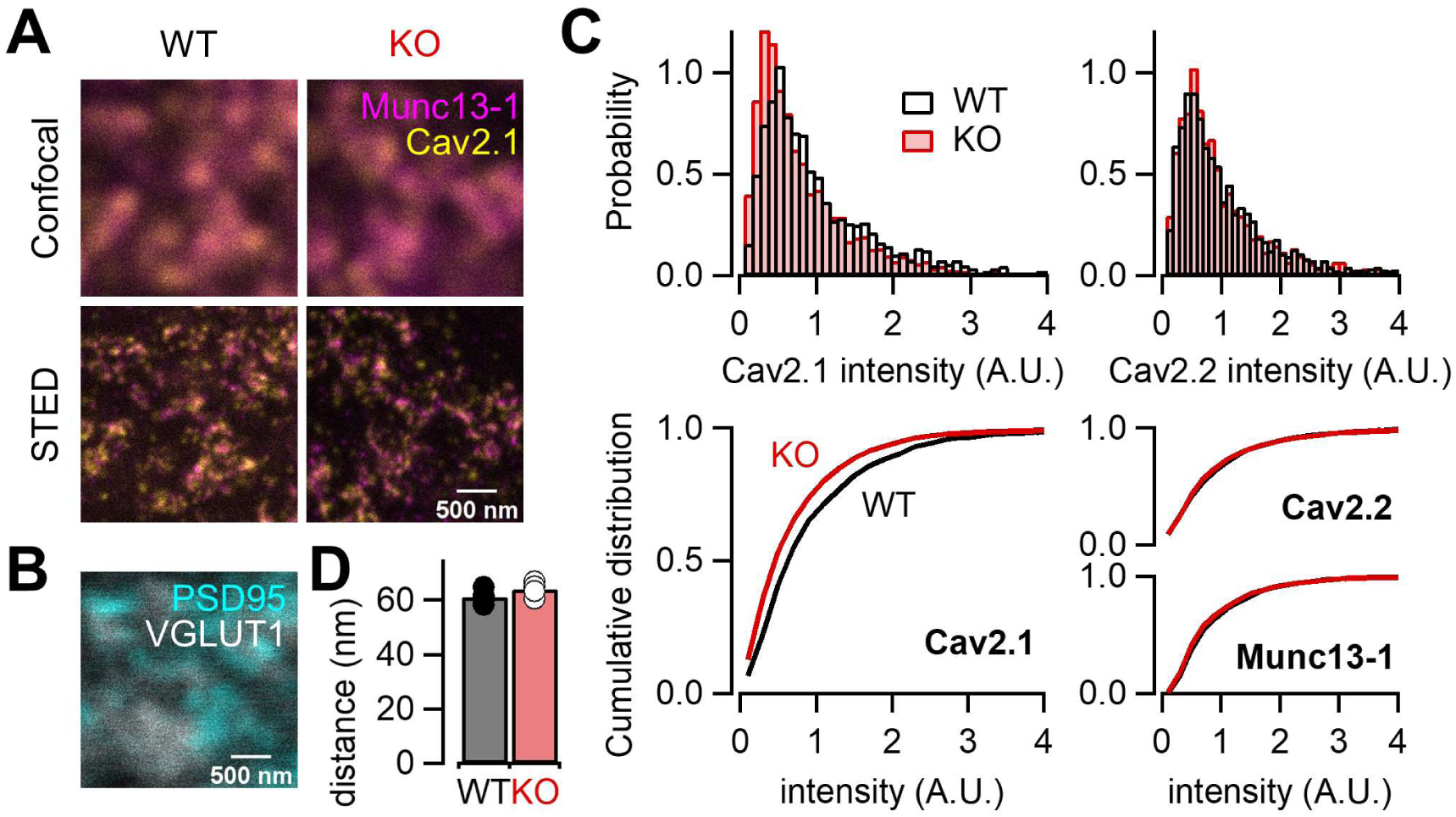
RIM-BP2 deletion alters the signal intensity of Cav2.1 clusters within the AZ. (**A**) Confocal *(top)* and STED *(bottom)* images of Munc13-1 (magenta) and Cav2.1 (yellow) clusters at hippocampal mossy fiber terminals of WT *(left)* and RIM-BP2 KO *(right)* mice. (Scale bar: 500 nm.) (**B**) Confocal images of PSD95 (cyan) and VGLUT1 (white) to identify glutamatergic synapses in CA3 stratum lucidum: mossy fiber-CA3 synapses. (Scale bar: 500 nm.) The image was taken from same region shown in (**A**), *(left)*. (**C**) Histograms *(top)* and cumulative distribution plots *(bottom)* of the total signal intensity of Cav2.1 *(left)*, Cav2.2, and Munc13-1 *(right)* at AZs in WT (black) and RIM-BP2 KO (red) mice. (**D**) The average nearest neighbor distance between Cav2.1 and Munc13-1 clusters in WT (black; n = 4 animals) and RIM-BP2 KO (red; n = 4 animals) mice. Several hundreds of AZs per image were analyzed. Data show the average value of distance per animal and error bars represent SEM. Each data point indicates individual values. Numerical values of plots are provided in Figure 7-source data 1.

**Figure 7-source data 1.** Datasets of numerical values presented in Figure 7.

**Figure 7-figure supplement 1.**
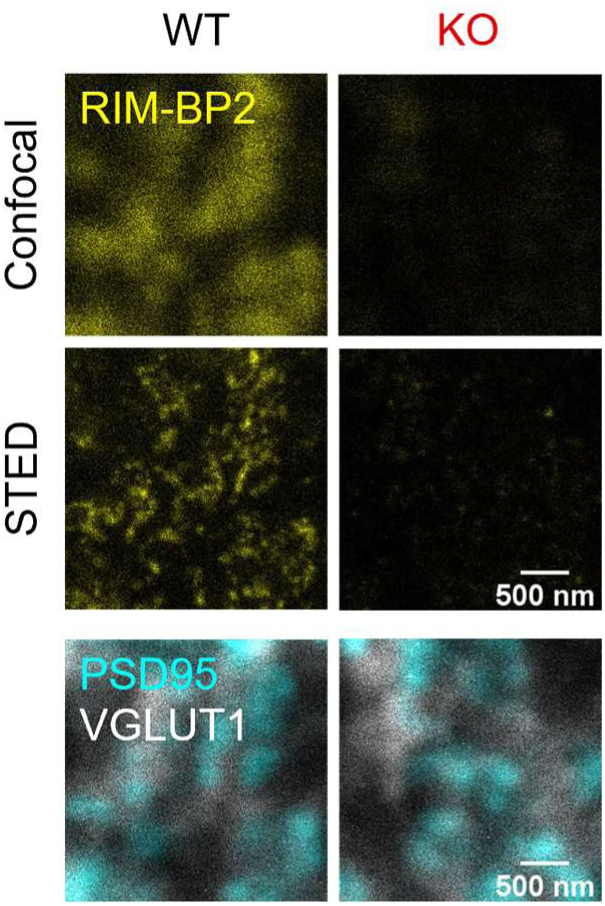
STED microscopy confirmed loss of RIM-BP2 proteins in KO terminals. Confocal *(top)* and STED *(middle)* images of RIM-BP2 (yellow) in WT *(left)* and RIM-BP2 KO *(right)* hippocampal mossy fiber terminals identified by PSD95 (cyan) and VGLUT1 (white) *(bottom)*. (Scale bar: 500 nm.) The *bottom* images show the same region as *top* and *middle*.

**Figure 7-figure supplement 2.**
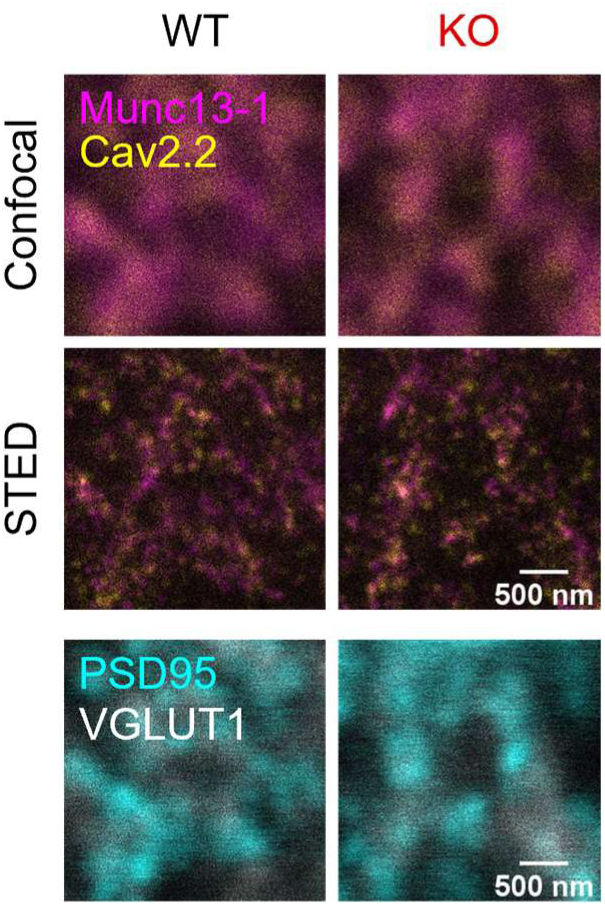
STED imaging of Cav2.2 at hippocampal mossy fiber terminals. Confocal *(top)* and STED *(middle)* images of Munc13-1 (magenta) and Cav2.2 (yellow) clusters in WT *(left)* and RIM-BP2 KO *(right)* mice. Confocal images of PSD95 (cyan) and VGLUT1 (white) in the same region as *top* and *middle* are also shown *(bottom)*. (Scale bar: 500 nm.)

**Figure 7-figure supplement 3.**
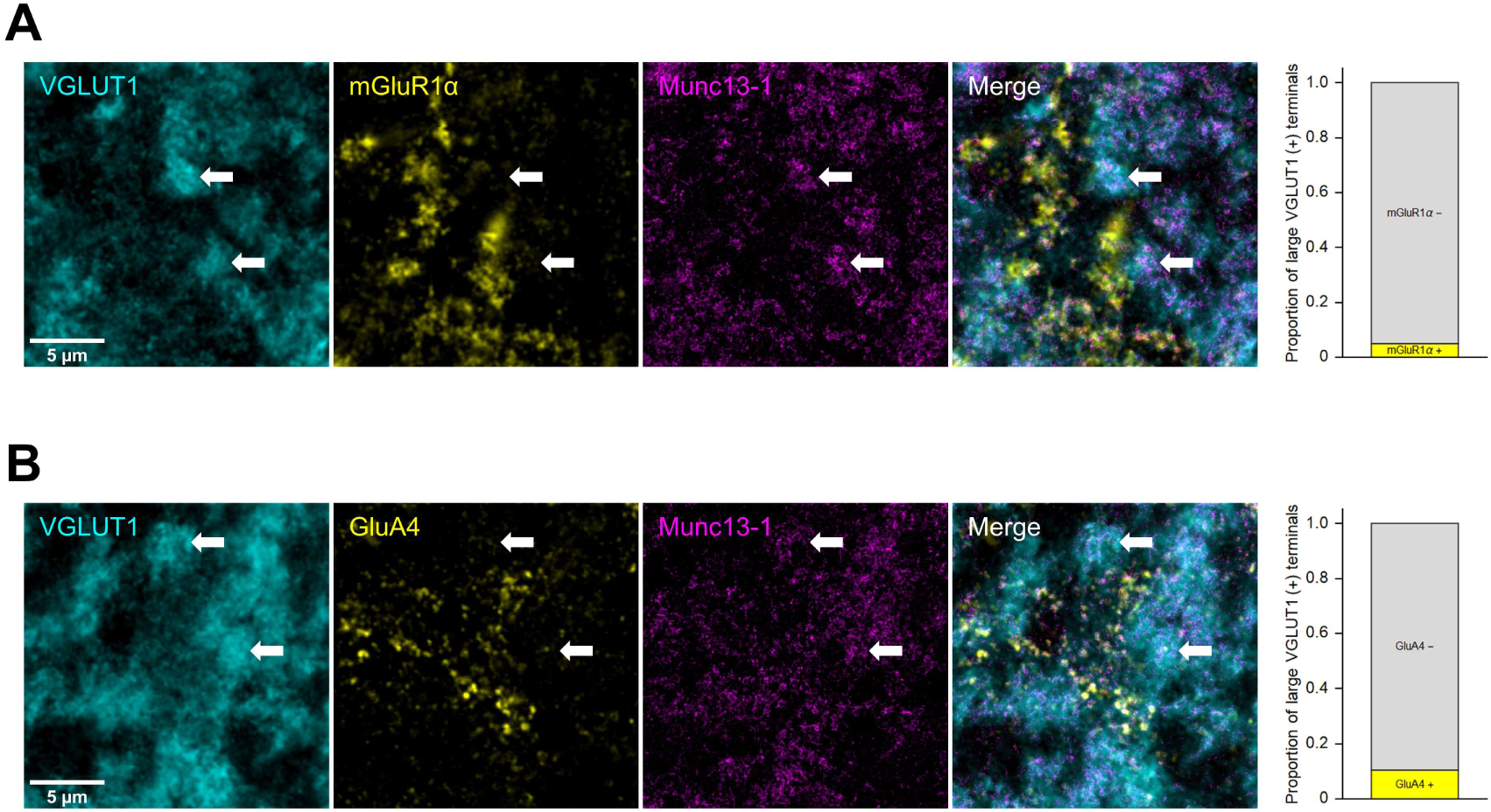
Immunohistochemical characterization of large VGLUT1 positive terminals in the CA3 stratum lucidum. **(A)** Confocal images of VGLUT1 (cyan) and mGluR1α (yellow) were taken with STED images of Munc13-1 (magenta) in the CA3 stratum lucidum. Arrows indicate representative large MF terminals identified by VGLUT1 signals (Scale bar: 5 µm.) The right graph shows the proportion of large VGLUT1 positive terminals contacting mGluR1α positive processes. (**B**) Same as (**A**) but for GluA4 instead of mGluR1α. (Scale bar: 5 µm.)

**Figure 7-figure supplement 4.**
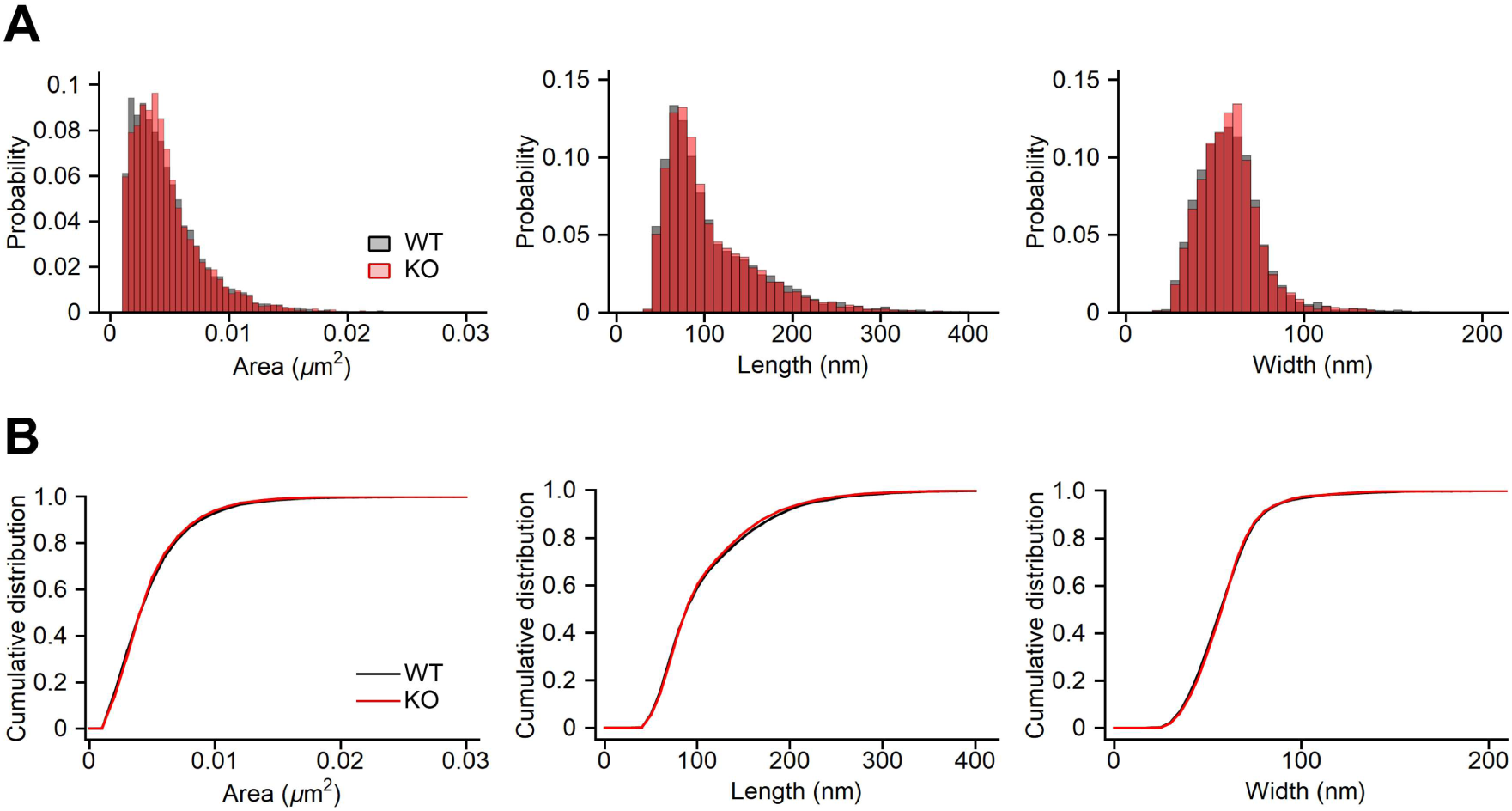
The area, length, width of Cav2.1 cluster. In A, distribution plots of area (left), length (middle) and width (right) are shown. In B, cumulative distribution plots are shown. The same samples as Fig. 7. Numerical values of plots are provided in Figure 7-figure supplement 4-source data 1.

**Figure 7-figure supplement 4-source data 1.** Datasets of numerical values presented in Figure 7-figure supplement 4.

**Figure 7-figure supplement 5.**
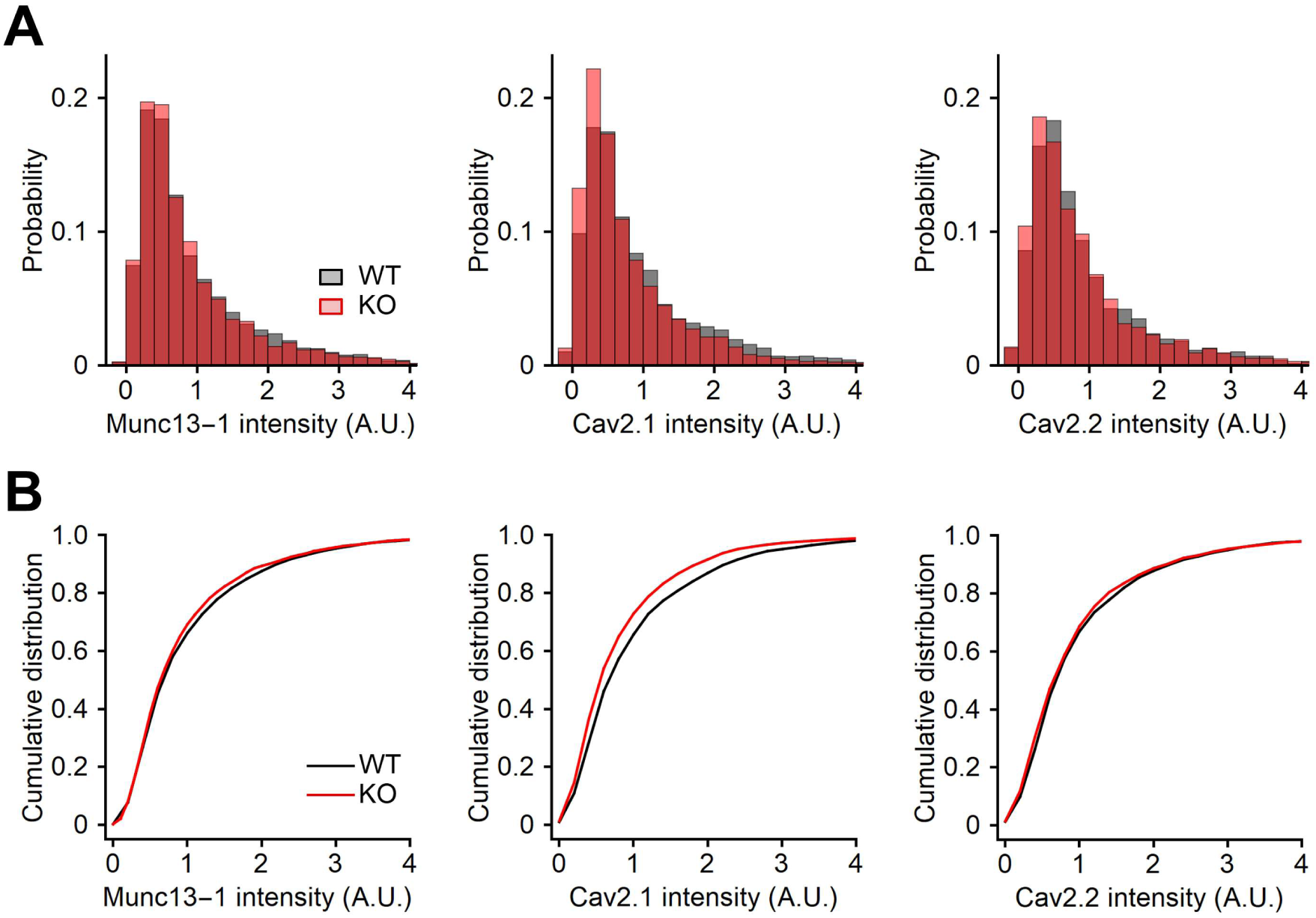
Quantification of the signal intensity using confocal images. The signal intensity of Munc13-1 *(left)*, Cav2.1 *(middle)*, and Cav2.2 *(right)* at AZs in WT (black) and RIM-BP2 KO (red) mice were quantified from confocal images. The position of AZ was determined from the Munc13-1 STED data, and the signal intensity of Munc13-1, Cav2.1 and Cav2.2 was measured. Histograms (**A**) and cumulative distribution plots (**B**) are shown. The same samples as Fig. 7. Numerical values of plots are provided in Figure 7-figure supplement 5-source data 1.

**Figure 7-figure supplement 5-source data 1.** Datasets of numerical values presented in Figure 7-figure supplement 5.

## Discussion

Accumulation of Ca^2+^ channels at the AZ, and efficient vesicle docking and priming are essential factors for fast synaptic vesicle exocytosis. AZ proteins are critical for fast synchronous release and synaptic diversity. RIM-BPs have been implicated in recruitment of Ca^2+^ channels to the AZ (Kaeser et al., 2011; Liu et al., 2011) by interacting with RIMs and Ca^2+^ channels (Wang et al., 2000; Hibino et al., 2002). In addition, RIM-BPs, like RIM1/2, recruit Munc-13 and accelerate synaptic vesicle priming (Brockmann et al., 2020). At murine central synapses, RIM-BPs deficient synapses demonstrate overall a rather mild decrease in the amounts of transmitter release (Acuna et al., 2015; Grauel et al., 2016; Luo et al., 2017; Krinner et al., 2017; Brockmann et al., 2019; Butola et al., 2021). Most studies so far have been mainly focused on synapses with high release probability (phasic synapses) such as the calyx of Held and CA3-CA1 synapses. That said, at mossy fiber-CA3 synapses, KO of RIM-BPs has a strong influence on synaptic transmission (Brockmann et al., 2019). Notably, at mossy fiber-CA3 synapses, release probability is relatively low (tonic synapses). It was unknown whether the reduction of transmitter release at RIM-BP2 KO mossy fiber synapses was due to impaired Ca^2+^ channel recruitment, and / or impaired vesicle docking or priming. Using STED microscopy, Brockmann et al. (2019) suggested that the physical recruitment of Munc13-1 might be an important function of RIM-BP2, though functional demonstration had been lacking.

By using direct presynaptic patch clamp recordings, we here observed a decrease of Ca^2+^ current amplitudes (∼30%) in RIM-BP2 KO mice (Fig. 1). Consistently, STED microscopy supported reduced abundance of P/Q-type Ca^2+^ channels (Cav2.1) in the mutant mossy fiber terminal (Fig. 7). Interestingly, this observation is similar to that at Drosophila NMJ and hair cell synapses (Liu et al., 2011; Krinner et al., 2017), but not that at other synapses (Acuna et al., 2015; Grauel et al., 2016; Butola et al., 2021), suggesting that the functional role of RIM-BP2 in recruiting Ca^2+^ channels differs among synapse types.

Brockmann et al. (2019) observed a reduction of Munc13-1 cluster number and docked vesicles in RIM-BP2 deficient synapses at hippocampal mossy fiber terminals (Fig. 7). They identified the terminals by using ZnT3, enriched at mossy fiber (Brockmann et al., 2019; Wenzel et al., 1997). Our STED imaging did not detect differences in Munc13-1 cluster number at the AZ between WT and RIM-BP2 KO mice. Notably, we analyzed the number of Cav2.1, Cav2.2 and Munc13-1 clusters only within AZs showing direct co-localization with VGLUT1 and PSD-95, which mainly reflects the AZs facing CA3 pyramidal cells rather than the synapses onto interneurons (Acsády et al., 1998; Rollenhagen et al., 2007). Such a difference in areas analyzed between previous and our study might explain the difference. In addition, some developmental variability/compensation could still occur in the range we studied and be relevant for the discrepancy between Munc13-1 results between Brockmann et al. (2019) and this study. Also, the spatial resolution of our study is limited and we cannot rigidly measure the signals only from large mossy fiber terminals. It is important to state that our results do not exclude roles of RIM-BP2 in synaptic vesicle priming. Indeed, we have shown that the reduction of the EPSC amplitudes could not be fully explained by the reduced Ca^2+^ influx (Fig. 5). Because the differences in the RRP size measured from capacitance measurements were relatively minor (Fig. 1-Fig. 3), the fusion competence was likely to be impaired by RIM-BP2 KO, but this could be overcome by large Ca^2+^ influx. Consistently, the EPSCs responses of KO became larger and comparable to those of WT (Fig. 6A, Fig. 6B, and Figure 6-figure supplement 1) during repetitive AP stimulation, at least qualitatively.

Recent studies have reported that RIM-BPs deletion altered Ca^2+^ channel localization and loosened the coupling between Ca^2+^ channels and synaptic vesicles at mammalian synapses (Acuna et al., 2015; Grauel et al., 2016; Butola et al., 2021). We investigated the Ca^2+^ channel-vesicle coupling at hippocampal mossy fiber terminals. The sensitivity of release to Ca^2+^ chelator EGTA was not changed between WT and RIM-BP2 KO mice. We also indicated RIM-BP2 KO did not alter the coupling of Ca^2+^ channels to release sites by using STED microscopy, though spatial resolution of STED microscopy in this study could not allow to detect small changes (Fig. 7).

At some mammalian synapses, deletion of RIM-BP2 affects physical distance between Ca^2+^ channels and synaptic vesicles (Acuna et al., 2015; Grauel et al., 2016; Butola et al., 2021). In contrast, at hippocampal mossy fiber terminals having low release probability (Vyleta and Jonas, 2014), RIM-BP2 KO decreased the abundance of P/Q-type Ca^2+^ channels, thereby reducing Ca^2+^ influx and neurotransmitter release. We hypothesize the specific molecular-architectural and biochemical contributions by RIM-BPs might be of different relevance among different types of synapses. Brockmann et al. (2020) proposed that RIM-BP2 controlled Ca^2+^ channel recruitment and vesicle priming by profound interaction with RIMs and Munc13s. These results have been obtained through studying hippocampal cultures dominated by phasic synapses. It seems possible that the interaction among three protein types is tight at some types of phasic synapses, simply because of high density of these proteins at AZs. Alternatively, one might speculate about additional proteins, different interaction surfaces taken, transsynaptic columns (Tang et al., 2016). Deletion of RIM-BP2 does not lead to loss of Ca^2+^ channels themselves and/or vesicle priming due to remaining RIMs, Munc13, and other proteins, which can regulate recruitment of Ca^2+^ channels and synaptic vesicle priming without RIM-BP2. Instead, fine-tuning of release such as synchronization is compromised. We hypothesize that AZ scaffold might be differently built, leading to different consequences when eliminating one component. Hence deletion of RIM-BP2 leads to immediate loss of Ca^2+^ channels and vesicle priming. This is in line with the proposal that at hippocampal mossy fiber synapses, synaptic vesicles are only loosely coupled with Ca^2+^ channels (Vyleta and Jonas, 2014) and vesicles are de-primed under resting conditions (Miki et al., 2016; Neher and Brose, 2018). Future research along this line may lead to an understanding of how fine molecular differences in the AZ scaffolds might orchestrate synaptic diversity.

Although this study reveals reduction of Ca^2+^ currents and reduced fusion competence in RIM-BP2 KO mossy fiber synapses, it has some limitation. Most importantly, with conventional knockouts the observed changes could reflect compensatory developmental changes and not direct outcomes of the RIM-BP2 knockout. This issue should be resolved by using region-specific and time-dependent conditional KO.

## Materials and Methods

### Slice preparation

All animal experiments were conducted in accordance with the guidelines of the Physiological Society of Japan, and were approved by Doshisha University Animal Experiment Committee (A22063, D22063).

We used male and female C57BL/6 mice (postnatal days 35-40). WT and RIM-BP2 KO mice (Grauel et al., 2016) were deeply anesthetized with isoflurane and decapitated. Their brains were quickly removed and chilled in sherbet-like cutting solution containing (in mM): 87 NaCl, 75 sucrose, 25 NaHCO_3_, 1.25 NaH_2_PO_4_, 2.5 KCl, 10 glucose, 0.5 CaCl_2_ and 7 MgCl_2_ bubbled with 95% O_2_ and 5% CO_2_ (Hallermann et al., 2003). Hippocampal slices (300 μm thick) were prepared from brains using a vibratome (VT1200S, Leica) in ice-cold cutting solution. After slicing, slices were incubated in cutting solution at 37°C for 30 min, and subsequently kept at room temperature (22-25°C) up to 4 h.

### Whole-cell recordings

Electrophysiological recordings were performed in a recording chamber filled with the extracellular solution containing (in mM): 125 NaCl, 2.5 KCl, 25 glucose, 25 NaHCO_3_, 1.25 NaH_2_PO_4_, 0.4 ascorbic acid, 3 myo-inositol, 2 Na-pyruvate, 2 CaCl_2_ and 1 MgCl_2_ saturated with 95% O_2_ and 5% CO_2_. In some experiments, the concentration of CaCl_2_ was changed to 1 mM, 1.5 mM and 4 mM. For presynaptic membrane capacitance measurements, 1 μM tetrodotoxin (TTX, Wako) was added to block Na^+^ channels. Moreover, for recording Ca^2+^ currents (Fig. 1B), 10 mM TEA-Cl was added to block K^+^ channels. Slices were visualized by an upright microscope (BX-51, Olympus). Whole-cell patch-clamp recordings were applied to hippocampal mossy fiber terminals and CA3 pyramidal cells at room temperature. The patch pipettes were filled with the intracellular solution containing (in mM): 135 Cs-gluconate, 20 TEA-Cl, 10 Hepes, 5 Na_2_-phosphocreatine, 4 MgATP, 0.3 GTP and 0.5 EGTA (pH = 7.2 adjusted with CsOH). In some experiments, the concentration of EGTA was changed to 5 mM. Presynaptic patch pipettes (BF150-86-10, Sutter Instrument) had a resistance of 15-20 MΩ and series resistance (R_s_) was 30-70 MΩ. R_s_ was compensated so that residual resistance was 30-35 MΩ. For postsynaptic recordings, the same internal solution was used, except that EGTA concentration was raised to 5 mM. Postsynaptic patch pipettes (BF150-86-10, Sutter Instrument) had a resistance of 3-7 MΩ and series resistance (R_s_) was 5-20 MΩ which was compensated by 50-80 %.

Patch-clamp recordings were performed using an EPC10/3 or EPC10/2 amplifier (HEKA) in voltage-clamp mode, controlled by Patchmaster software (HEKA). Membrane currents were low-pass filtered at 2.9 kHz and sampled at 20 or 50 kHz.

Membrane capacitance measurements were performed using an EPC10/3 amplifier in the sine + DC configuration (Gillis, 2000) using Patchmaster software (HEKA). A sine wave (30 mV in amplitude, 1000 Hz in frequency) was superimposed on the holding potential of -80 mV. The terminal was depolarized from -80 mV to +10 mV for 5-100 ms. The IV of presynaptic Ca currents were measured by sequential depolarized for 5 ms, with 2 ms intervals from -80 mV to +70 mV by 10 mV steps.

For the EPSC measurements, postsynaptic cells (CA3 pyramidal cells) were voltage clamped at -60 - -80 mV. For Figs. 5 and 6, the data were corrected for the different holding potential (re-calculated to the value at -70 mV assuming the reversal potential of 0 mV, which may have the correction effect of ∼15 %). At the end of the experiments, DCG-Ⅳ was applied to confirm mossy fiber synaptic responses, whenever possible. The mossy fibers were stimulated extracellularly using a glass electrode used for patch clamp containing the extracellular solution and the stimulation intensity was usually 10-60 V.

Data were obtained from at least three different WT and RIM-BP2 KO terminals in each set of experiments (biological replicate).

### Immunostaining

Brains of WT and RIM-BP2 KO mice were quickly removed and transferred to cryomold (Sakura Finetek) filled with tissue freezing medium (Leica). Brains were instantaneously frozen on aluminum block in liquid nitrogen and stored at -80°C. For cryosectioning, frozen brains were kept in cryostat (CM1860, Leica) at -18°C for 30 min. Then, 10 μm-thick sections were sliced and collected on cover glasses (Matsunami glass, 25×25 No.1). Sections were quickly fixed by dehydration with a heat blower at 50°C for 1 min, and further dehydrated in ethanol for 30 min at -25°C and in acetone for 10 min on ice. After blocking with PBS containing 0.3% BSA at room temperature, sections were incubated with primary antibodies diluted in PBS containing 0.3% BSA: anti-VGLUT1 (1:2000, Synaptic Systems, guinea pig polyclonal, RRID:<u>AB_887878</u>), anti-PSD-95 (1:50, NeuroMab, mouse monoclonal IgG2a, clone K28/43, TC Supernatant, RRID:<u>AB_2877189</u>), anti-Munc13-1 (1:1000, mouse monoclonal IgG1, clone 11B-10G, Sakamoto et al., 2018), and anti-Cav2.1 (1:400, Synaptic Systems, rabbit polyclonal, RRID:<u>AB_2619841</u>) or anti-Cav2.2 (1:400, Synaptic Systems, mouse monoclonal IgG2b, clone 163E3, RRID:<u>AB_2619843</u>) or anti-RIM-BP2 (1:200, mouse monoclonal IgG2b, clone 8-4G, Sakamoto et al., 2022) for 3 hours at room temperature. Afterwards, sections were washed and incubated with fluorescence dye labeled secondary antibodies: DyLight 405 Anti-Guinea Pig IgG (Jackson ImmunoResearch, RRID:<u>AB_2340470</u>), Alexa Fluor 488 (Thermo Fisher Scientific) labeled Anti-Mouse IgG2a (Jackson ImmunoResearch, RRID:<u>AB_2338462</u>), STAR635P (Abberior) labeled Anti-Mouse IgG1 (Jackson ImmunoResearch, RRID:<u>AB_2338461</u>), and Alexa Fluor 594 (Thermo Fisher Scientific) labeled Anti-Rabbit IgG (Jackson ImmunoResearch, RRID:<u>AB_2340585</u>) or Anti-Mouse IgG2b (Jackson ImmunoResearch, RRID:<u>AB_2338463</u>) for 1 hour at room temperature. Sections were post-fixed with 4% PFA in PBS and washed with PBS. Sections were mounted on coverslips using Prolong Glass Antifade Mountant (Thermo Fisher Scientific). The same number of sections were obtained from WT and RIM-BP2 KO brains and processed in parallel at the same experimental day (biological replicate).

For Figure 7-figure supplement 3, only WT brains were used. Primary antibodies used were anti-VGLUT1 (1:2000, Synaptic Systems, rabbit polyclonal, RRID:AB_887877), anti-VGLUT1 (1:2000, Synaptic Systems, guinea pig polyclonal, RRID:<u>AB_887878</u>), anti-mGluR1α (1:200, Nittobo Medical, guinea pig polyclonal, mGluR1a-GP-Af660), anti-GluA4C (1:100, Nittobo Medical, rabbit polyclonal, GluR4C-Rb-Af160), and anti-Munc13-1 (1:1000, mouse monoclonal IgG1, clone 11B-10G).

### STED imaging

STED imaging was performed using TCS SP8 STED 3x microscope (Leica) equipped with a 405 nm diode laser, a pulsed white light laser (WLL), a continuous 592 nm STED laser for alignment, a pulsed 775 nm STED laser, HyD detectors, and a 100× oil-immersion objective lens (NA = 1.4). STED images were acquired using the Leica LAS-X software with an image format of 1024×1024 pixels, 16-bit sampling, 8-line accumulations, and 11.36-zoom factor, yielding a pixel size of 10 nm. HyD detectors were configured to counting mode with a gating from 0.5 to 6.5 nanosecond. A 405 nm diode laser was used to excite DyLight405. Alexa488, Alexa594, and STAR635P were excited using WLL at 488 nm, 561 nm, and 633 nm, respectively. The 775 nm STED laser power was set to 75% and 100% of maximum power for depletion of STAR635P and Alexa594, respectively, and delay time was set to 300 picoseconds. STED imaging was performed on thin sections obtained from four different WT and RIM-BP2 KO mice (biological replicate). Images were acquired three times from each section (technical replicate).

### Analysis

For electrophysiological experiments, the data were analyzed by Igor Pro (WaveMetrics) or Excel (Microsoft Corp). Presynaptic calcium currents were measured at the peak amplitude during the depolarizing pulse. Capacitance jumps were measured between baseline (before the pulse) and ∼10 ms after the pulse where the effect of tail currents became minor. To determine the time constant of release, an exponential fit was applied. Values are given as mean ± SEM, and n indicates the number of recorded terminals or the number of animals used. Statistical method of sample size determination was not done, but our sample sizes are similar to those of previous studies (Hallermann et al., 2003, Midorikawa and Sakaba, 2017).

For STED imaging, data were analyzed using custom-designed programs in Mathematica (Wolfram), values were processed by Igor Pro (WaveMetrics), and pictures were created with ImageJ (NIH). To determine the area of AZs, image masks were generated from STED images of Munc13-1 by unsharp masking and image binarization. Only AZs that show co-localization with VGLUT1 and PSD-95 immunofluorescence signals were included in the analysis. To confine the analysis to the large mossy fiber boutons, AZs in presynaptic terminals of small VGLUT1-positive area were excluded. Then, the background subtracted integral of signal intensity of Cav2.1 or Cav2.2 was quantified using the image masks of AZs. The background signal intensity was estimated from non-AZ area in the same STED images. For quantification of Cav2.1 clusters in the AZs, STED images of Cav2.1 were deconvolved using a gaussian kernel with a radius of 40 nm. Image masks of each Ca^2+^ channel were generated from deconvolved STED images by unsharp masking and image binarization. Then, the total signal intensity, the size, and the number of Cav2.1 clusters at the AZs were quantified using the image masks of Cav2.1 clusters. Custom-designed programs in Mathematica (Wolfram) are available from Source code 1. Values are given as mean ± SEM, and n indicates the number of animals. Statistical method of sample size determination was not done, but our sample sizes are similar to those of previous studies (Brockmann et al., 2019).

Experiments were not fully randomized and blinded.

Statistical analysis was done in MATLAB (The MathWorks) or SPSS (IBM). t-test, ANOVA, and mixed model were used for statistical tests. Significance level was set at α = 0.05, and p values were described in the main text or figure legends.

## Data availability

All data generated in this study are included in the main text or shown as scatter plots in each graph. Numerical values of graphs are provided in Source data files. The custom-written code files in Mathematica are uploaded as Source code file.

## Acknowledgements

We thank Stephan Sigrist, Dietmar Schmitz and Christian Rosenmund for kindly providing us RIM-BP2 KO mice. We also thank Stephan Sigrist for comments on the manuscript and The IRCN Imaging Core, The University of Tokyo Institutes for Advanced Studies, for the use of STED microscopy and for assistance. This works has been supported by JSPS KAKENHI (JP20J20550 to RM, JP20KK0171, JP21K15183 to HS, JP20H03427 to KH, JP20KK0171, JP21H02598 to TS), JSPS Core-to-Core program A. Advanced Research Networks (JPJSCCA20220007 to TS), JST PRESTO (JPMJPR21E7 to HS) and Takeda Science Foundation (Bioscience and Specific Research Grants to TS).

## Competing interests

None

## Notes

### Competing Interest Statement

The authors have declared no competing interest.

### Summary of Updates

Figure 7-figure supplement 5 was added. Some small textual changes.

## References

Acsády L, Kamondi A, Sík A, Freund T, Buzsáki G. 1998. GABAergic cells are the major postsynaptic targets of mossy fibers in the rat hippocampus. Journal of Neuroscience 18:3386–3403. DOI: 10.1523/JNEUROSCI.18-09-03386.1998, PMID: 9547246

Acuna C, Liu X, Gonzalez A, Südhof TC. 2015. RIM-BPs Mediate Tight Coupling of Action Potentials to Ca^2+^-Triggered Neurotransmitter Release. Neuron 87:1234–1247. DOI: 10.1016/j.neuron.2015.08.027, PMID: 26402606

Adler EM, Augustine GJ, Duffy SN, Charlton MP. 1991. Alien intracellular calcium chelators attenuate neurotransmitter release at the squid giant synapse. Journal of Neuroscience 11:1496–1507. DOI: 10.1523/JNEUROSCI.11-06-01496.1991, PMID: 1675264

Blatow M, Caputi A, Burnashev N, Monyer H, Rozov A. 2003 Ca^2+^ buffer saturation underlies paired pulse facilitation in calbindin-D28k-containing terminals. Neuron 38: 79–88. DOI: 10.1016/s0896-6273(03)00196-x, PMID: 12691666

Bollmann JH, Sakmann B, Borst JG. 2000. Calcium sensitivity of glutamate release in a calyx-type terminal. Science 289: 953–957. DOI: 10.1126/science.289.5481.953, PMID: 10937999

Borst JG, Sakmann B. 1996. Calcium influx and transmitter release in a fast CNS synapse. Nature 383:431–434. DOI: 10.1038/383431a0, PMID: 8837774

Brockmann MM, Maglione M, Willmes CG, Stumpf A, Bouazza BA, Velasquez LM, Grauel MK, Beed P, Lehmann M, Gimber N, Schmoranzer J, Sigrist SJ, Rosenmund C, Schmitz D. 2019. RIM-BP2 primes synaptic vesicles *via* recruitment of Munc13-1 at hippocampal mossy fiber synapses. eLife 8:e43243. DOI: 10.7554/eLife.43243, PMID: 31535974

Brockmann MM, Zarebidaki F, Camacho M, Grauel MK, Trimbuch T, Südhof TC, Rosenmund C. 2020. A Trio of Active Zone Proteins Comprised of RIM-BPs, RIMs, and Munc13s Governs Neurotransmitter Release. Cell Reports 32:107960-107960. DOI: 10.1016/j.celrep.2020.107960, PMID: 32755572

Butola T, Alvanos T, Hintze A, Koppensteiner P, Kleindienst D, Shigemoto R, Wichmann C, Moser T. 2021. RIM-Binding Protein 2 Organizes Ca^2+^ Channel Topography and Regulates Release Probability and Vesicle Replenishment at a Fast Central Synapse. Journal of Neuroscience 41:7742–7767. DOI: 10.1523/JNEUROSCI.0586-21.2021, PMID: 34353898

Castillo PE, Weisskopf MG, Nicoll RA. 1994. The role of Ca^2+^ channels in hippocampal mossy fiber synaptic transmission and long-term potentiation. Neuron 12:261–269. DOI: 10.1016/0896-6273(94)90269-0, PMID: 8110457

Dodge FA, Rahamimoff R. 1967. Co-operative action a calcium ions in transmitter release at the neuromuscular junction. The Journal of physiology 193:419–432. DOI: 10.1113/jphysiol.1967.sp008367, PMID: 6065887

Geiger JR, Jonas P. 2000. Dynamic control of presynaptic Ca^2+^ inflow by fast-inactivating K^+^ channels in hippocampal mossy fiber boutons. Neuron 28: 927–939. DOI: 10.1016/s0896-6273(00)00164-1, PMID: 11163277

Grauel MK, Maglione M, Reddy-Alla S, Willmes CG, Brockmann MM, Trimbuch T, Rosenmund T, Pangalos M, Vardar G, Stumpf A, Walter AM, Rost BR, Eickholt BJ, Haucke V, Schmitz D, Sigrist SJ, Rosenmund C. 2016. RIM-binding protein 2 regulates release probability by fine-tuning calcium channel localization at murine hippocampal synapses. PNAS 113:11615–11620. DOI: 10.1073/pnas.1605256113, PMID: 27671655

Hallermann S, Pawlu C, Jonas P, Heckmann M. 2003. A large pool of releasable vesicles in a cortical glutamatergic synapse. PNAS 100:8975–8980. DOI: 10.1073/pnas.1432836100, PMID: 12815098

Hibino H, Pironkova R, Onwumere O, Vologodskaia M, Hudspeth AJ, Lesage F. 2002. RIM binding proteins (RBPs) couple Rab3-interacting molecules (RIMs) to voltage-gated Ca^2+^ channels. Neuron 34:411–423. DOI: 10.1016/s0896-6273(02)00667-0, PMID: 11988172

Kaeser PS, Deng L, Wang Y, Dulubova I, Liu X, Rizo J, Südhof TC. 2011. RIM proteins tether Ca^2+^ channels to presynaptic active zones via a direct PDZ-domain interaction. Cell 144:282–295. DOI: 10.1016/j.cell.2010.12.029, PMID: 21241895

Katz B. 1969. The release of neural transmitter substances. Liverpool University Press. ISBN: 0853230609

Krinner S, Butola T, Jung S, Wichmann C, Moser T. 2017. RIM-Binding Protein 2 Promotes a Large Number of Ca_V_1.3 Ca^2+^-Channels and Contributes to Fast Synaptic Vesicle Replenishment at Hair Cell Active Zones. Frontiers in Cellular Neuroscience 11:334. DOI: 10.3389/fncel.2017.00334, PMID: 29163046

Krinner S, Predoehl F, Burfeind D, Vogl C, Moser T. 2021. RIM-Binding Proteins Are Required for Normal Sound-Encoding at Afferent Inner Hair Cell Synapses. Frontiers in molecular neuroscience 14:651935. DOI: 10.3389/fnmol.2021.651935, PMID: 33867935

Li L, Bischofberger J, Jonas P. 2007. Differential gating and recruitment of P/Q-, N-, and R-type Ca^2^^+^ channels in hippocampal mossy fiber boutons. *Journal of Neuroscience* **27**:13420-13429. DOI: 10.1523/JNEUROSCI.1709-07.2007, PMID: 18057200

Lindau M, Neher E. 1988. Patch-clamp techniques for time-resolved capacitance measurements in single cells. Pflügers Archiv 411:137–146. DOI: 10.1007/BF00582306, PMID: 3357753

Liu KS, Siebert M, Mertel S, Knoche E, Wegener S, Wichmann C, Matkovic T, Muhammad K, Depner H, Mettke C, Bückers J, Hell SW, Müller M, Davis GW, Schmitz D, Sigrist SJ. 2011. RIM-binding protein, a central part of the active zone, is essential for neurotransmitter release. Science 334:1565–1569. DOI: 10.1126/science.1212991, PMID: 22174254

Luo F, Liu X, Südhof TC, Acuna C. 2017. Efficient stimulus-secretion coupling at ribbon synapses requires RIM-binding protein tethering of L-type Ca^2+^ channels. Proceedings of the National Academy of Sciences 114:E8081–E8090. DOI: 10.1073/pnas.1702991114, PMID: 28874522

Midorikawa M, Sakaba T. 2017. Kinetics of Releasable Synaptic Vesicles and Their Plastic Changes at Hippocampal Mossy Fiber Synapses. Neuron 96:1033–1040.e3. DOI: 10.1016/j.neuron.2017.10.016, PMID: 29103807

Miki T, Malagon G, Pulido C, Llano I, Neher E, Marty A. 2016. Actin- and Myosin-Dependent Vesicle Loading of Presynaptic Docking Sites Prior to Exocytosis. Neuron 91:808–823. DOI: 10.1016/j.neuron.2016.07.033, PMID: 27537485

Mittelstaedt T, Schoch S. 2007. Structure and evolution of RIM-BP genes: identification of a novel family member. Gene 403:70–79. DOI: 10.1016/j.gene.2007.08.004, PMID: 17855024

Müller M, Genç Ö, Davis GW. 2015. RIM-binding protein links synaptic homeostasis to the stabilization and replenishment of high release probability vesicles. Neuron 85:1056–1069. DOI: 10.1016/j.neuron.2015.01.024, PMID: 25704950

Neher E, Brose N. 2018. Dynamically Primed Synaptic Vesicle States: Key to Understand Synaptic Short-Term Plasticity. Neuron 100:1283–1291. DOI: 10.1016/j.neuron.2018.11.024, PMID: 30571941

Pelkey KA, Topolnik L, Lacaille J-C, McBain CJ. 2006. Compartmentalized Ca^2+^ channel regulation at divergent mossy-fiber release sites underlies target cell-dependent plasticity. Neuron 52:497–510. DOI: 10.1016/j.neuron.2006.08.032, PMID: 17088215

Petzoldt AG, Götz TWB, Driller JH, Lützkendorf J, Reddy-Alla S, Matkovic-Rachid T, Liu S, Knoche E, Mertel S, Ugorets V, Lehmann M, Ramesh N, Beuschel CB, Kuropka B, Freund C, Stelzl U, Loll B, Liu F, Wahl MC, Sigrist SJ. 2020. RIM-binding protein couples synaptic vesicle recruitment to release sites. Journal of Cell Biology 219:e201902059. DOI: 10.1083/jcb.201902059, PMID: 32369542

Rollenhagen A, Sätzler K, Rodríguez EP, Jonas P, Frotscher M, Lübke JHR. 2007. Structural determinants of transmission at large hippocampal mossy fiber synapses. Journal of Neuroscience 27:10434–10444. DOI: 10.1523/JNEUROSCI.1946-07.2007, PMID: 17898215

Sakamoto H, Ariyoshi T, Kimpara N, Sugao K, Taiko I, Takikawa K, Asanuma D, Namiki S, Hirose K. 2018. Synaptic weight set by Munc13-1 supramolecular assemblies. Nature neuroscience 21:41–49. DOI: 10.1038/s41593-017-0041-9, PMID: 29230050

Sakamoto H, Kimpara N, Namiki S, Hamada S, Ohtsuka T, Hirose K. 2022. Synapse type-specific molecular nanoconfigurations of the presynaptic active zone in the hippocampus identified by systematic nanoscopy [unpublished manuscript]. bioRxiv 2022.03.11.483942. DOI: 10.1101/2022.03.11.483942

Schneggenburger R, Meyer AC, Neher E. 1999. Released fraction and total size of a pool of immediately available transmitter quanta at a calyx synapse. Neuron 23:399–409. DOI: 10.1016/s0896-6273(00)80789-8, PMID: 10399944

Schneggenburger R, Neher E. 2000. Intracellular calcium dependence of transmitter release rates at a fast central synapse. Nature 406: 889–893. DOI: 10.1038/35022702, PMID: 10972290

Südhof TC. 2012. The presynaptic active zone. Neuron 75:11–25. DOI: 10.1016/j.neuron.2012.06.012, PMID: 22794257

Tang AH, Chen H, Li TP, Metzbower SR, MacGillavry HD, Blanpied TA. 2016. A trans-synaptic nanocolumn aligns neurotransmitter release to receptors. Nature 536:210–214. DOI: 10.1038/nature19058, PMID: 27462810

Thanawala MS, Regehr WG. 2013. Presynaptic calcium influx controls neurotransmitter release in part by regulating the effective size of the readily releasable pool. Journal of Neuroscience 33:4625–4633. DOI: 10.1523/JNEUROSCI.4031-12.2013, PMID: 23486937

von Gersdorff H, Matthews G. 1996. Calcium-dependent inactivation of calcium current in synaptic terminals of retinal bipolar neurons. Journal of Neuroscience 16:115–122. DOI: 10.1523/JNEUROSCI.16-01-00115.1996, PMID: 8613777

Vyleta NP, Jonas P. 2014. Loose coupling between Ca^2+^ channels and release sensors at a plastic hippocampal synapse. Science 343:665–670. DOI: 10.1126/science.1244811, PMID: 24503854

Wadel K, Neher E, Sakaba T. 2007. The coupling between synaptic vesicles and Ca^2+^ channels determines fast neurotransmitter release. Neuron 53:563–575. DOI: 10.1016/j.neuron.2007.01.021, PMID: 17296557

Wang Y, Sugita S, Südhof TC. 2000. The RIM/NIM family of neuronal C2 domain proteins. Interactions with Rab3 and a new class of Src homology 3 domain proteins. Journal of Biological Chemistry 275:20033-20044. DOI: 10.1074/jbc.M909008199, PMID: 10748113

Wenzel HJ, Cole TB, Born DE, Schwartzkroin PA, Palmiter RD. 1997. Ultrastructural localization of zinc transporter-3 (ZnT-3) to synaptic vesicle membranes within mossy fiber boutons in the hippocampus of mouse and monkey. PNAS 94:12676–12681. DOI: 10.1073/pnas.94.23.12676, PMID: 9356509

